# A unique mineralizing pool of Gli1+ stem cells builds the tendon enthesis and demonstrates therapeutic potential

**DOI:** 10.1101/2022.02.17.480929

**Authors:** Fei Fang, Yang Xiao, Elazar Zelzer, Kam W. Leong, Stavros Thomopoulos

## Abstract

The enthesis, a fibrocartilaginous transition between tendon and bone, is necessary for the transfer of force from muscle to bone to produce joint motion. The enthesis is prone to injury due to mechanical demands, and it cannot regenerate. A better understanding of how the enthesis develops will lead to more effective therapies to prevent pathology and promote regeneration. Here, we used single-cell RNA sequencing to define the development transcriptome of the entheses over postnatal stages. Six resident cell types, including enthesis progenitors and mineralizing chondrocytes, were identified along with their transcription factor regulons and temporal regulation. Following our prior discovery of the necessity of Gli1-lineage cells for enthesis development and healing, we then examined their transcriptomes at single cell resolution and demonstrated their clonogenicity and multipotency. Transplantation of these Gli1- lineage cells to enthesis injuries improved healing, demonstrating their therapeutic potential for enthesis regeneration.

**Highlights:**

- The transcriptome and differentiation trajectory of enthesis stem cells during postnatal development are defined at single cell resolution.
- Transcription factor regulons drive enthesis stem cell differentiation.
- Gli1-lineage enthesis stem cells demonstrate *in vivo* and *in vitro* clonogenicity and multipotency.
- Transplantation of Gli1-lineage enthesis stem cells to enthesis injuries improves healing.

## Introduction

Tendon connects to bone via a specialized interface known as the enthesis (Lu and Thomopoulos, 2013). Enthesis disorders, including rotator cuff disease, psoriatic arthritis, and spondyloarthritis, are prevalent and result in a heavy clinical burden (Derwin et al., 2018; Schett et al., 2017). The healthy enthesis is a functionally graded tissue that bridges unmineralized tendon and mineralized bone, and is formed and maintained by a spatially graded distribution of cell phenotypes (Genin et al., 2009; Thomopoulos et al., 2010). The functionally graded tissue, necessary for reducing stress concentrations at the interface between two dissimilar materials, is not regenerated after enthesis degeneration or injury, resulting in a mechanically weak attachment and ultimately rupture (Carpenter et al., 1998; Thomopoulos et al., 2003). For this reason, failure rates after surgical repair of torn tendon to bone are remarkably high (Galatz et al., 2004; Harryman et al., 2003). Despite significant treatment advances over the past decade, the development of therapeutic strategies for regenerating the enthesis has been hampered by an incomplete understanding of enthesis development. In particular, it remains unclear how a spatial gradient in cell phenotypes forms across the enthesis to drive the formation of a mineral gradient from tendon to bone. With the maturity of single-cell RNA sequencing (scRNA-seq) technology, it is now possible to evaluate the transcriptional signatures of the cell phenotypes across the enthesis at single-cell resolution.

Resident stem cells have been identified for bone (i.e., bone marrow-derived mesenchymal stem cells) and tendon, but not for the enthesis. Bone marrow-derived mesenchymal stem cells have been studied for many decades (Alvarez-Dolado et al., 2003; Chan et al., 2018; Graham et al., 1990; Jiang et al., 2002). More recently, tendon stem cells have been isolated and characterized, demonstrating typical stem cell features, including self-renewal capacity, clonogenicity, and multipotency (Bi et al., 2007). Due to a lack of distinct and reliable markers to identify tendon stem cells, their *in vivo* identity, contribution to tendon pathology, and translational application remain elusive (Millar et al., 2021). Recently, cells expressing Gli1 (glioma-associated oncogene homolog1, indicative of activated hedgehog signaling), have been proposed to function as progenitors residing in the tendon enthesis and in the epitenon (Felsenthal et al., 2018; Wang et al., 2017b). Our previous lineage-tracing experiments demonstrated that Gli1-lineage (Gli1+) cells build the tendon enthesis and their ablation at early postnatal timepoints leads to defects in enthesis mineralization and biomechanical function (Schwartz et al., 2015). Furthermore, neonatal enthesis injuries showed excellent capacity to heal, and this process involved Gli1+ enthesis cells from the early postnatal period (Schwartz et al., 2017). Consistent with these findings, multiple studies have also demonstrated that Gli1+ cells form an essential niche in the colon by renewing stem cells, drive bone marrow fibrosis by activating cell expansion and myofibroblast differentiation, and regulate bone formation and fracture repair by differentiating into chondrocytes and osteoblasts (Degirmenci et al., 2018; Kramann et al., 2015; Schneider et al., 2017; Shi et al., 2017; Zhao et al., 2014). Gli1 therefore serves as a putative stem cell marker with particular relevance to the enthesis. However, the characteristics of enthesis Gli1 stem cells and their therapeutic potential have not been explored.

Herein, we revealed enthesis cell heterogeneity and identified six cell sub-populations using scRNA-seq. We used two different algorithms for scRNA-seq analysis to infer cell differentiation trajectories for enthesis stem cells differentiating into mineralizing chondrocytes. A gene regulatory network analysis and fluorescent *in situ* hybridization (FISH) were then used to identify a number of transcription factors coordinating tenogenesis, chondrogenesis, and osteogenesis to form an enthesis with spatially graded mineralization. To further define the enthesis stem cell population, enthesis Gli1+ cells were isolated and their transcriptomes were characterized at single cell resolution. These Gli1+ cells were shown to have colony formation capacity and multipotency. Finally, Gli1+ stem cells were transplanted into injured entheses and shown to promote extracellular matrix deposition and increased mineralization, demonstrating a promising therapeutic strategy for enthesis regeneration.

## Results

### Single-cell transcriptomes of enthesis mesenchymal cells demonstrate six distinct cell phenotypes

To capture single-cell transcriptomes of enthesis cells during the critical postnatal mineralization period, we isolated mouse supraspinatus tendon entheses from shoulders of C57BL/J6 mice at postnatal day 7 (P7), P18, and P56 and dissociated the enthesis cells for scRNA-seq analysis using the 10x Genomics platform (Figure S1A, B). After the elimination of doublets, dead, and apoptotic cells, 3,000-5,000 single cells with 61,990-106,444 reads/cell and 1,901-3,154 genes/cell for each timepoint passed quality control and were processed for downstream evaluation (Figure S1C). After data integration of different timepoints, dimensional reduction, and UMAP (uniform manifold approximation and projection) visualization using the Seurat V3 package, 7 distinct cell clusters were manually annotated according to differentially expressed genes and a set of marker genes (Figures 1A and S1D). Similarly, enthesis mesenchymal cells were selected and clustered with high resolution and cell subpopulations were annotated as enthesis progenitors, pre-enthesoblasts, enthesoblasts, mineralizing chondrocytes, tenoblasts/osteoblasts, and osteocytes (Figure 1B). Strikingly, the greatest percentage of mineralizing chondrocytes (normalized by the number of enthesis stromal cells) was observed at P18 compared to P11 and P56 (Figure 1C). This observation is consistent with previous research, which revealed active enthesis mineralization between P14 and P21 (Fang et al., 2020; Schwartz et al., 2015; Schwartz et al., 2012). In contrast, the proportion of the other cell subpopulations remained steady across developmental stages (Figures 1C and S2A).

**Figure 1.**
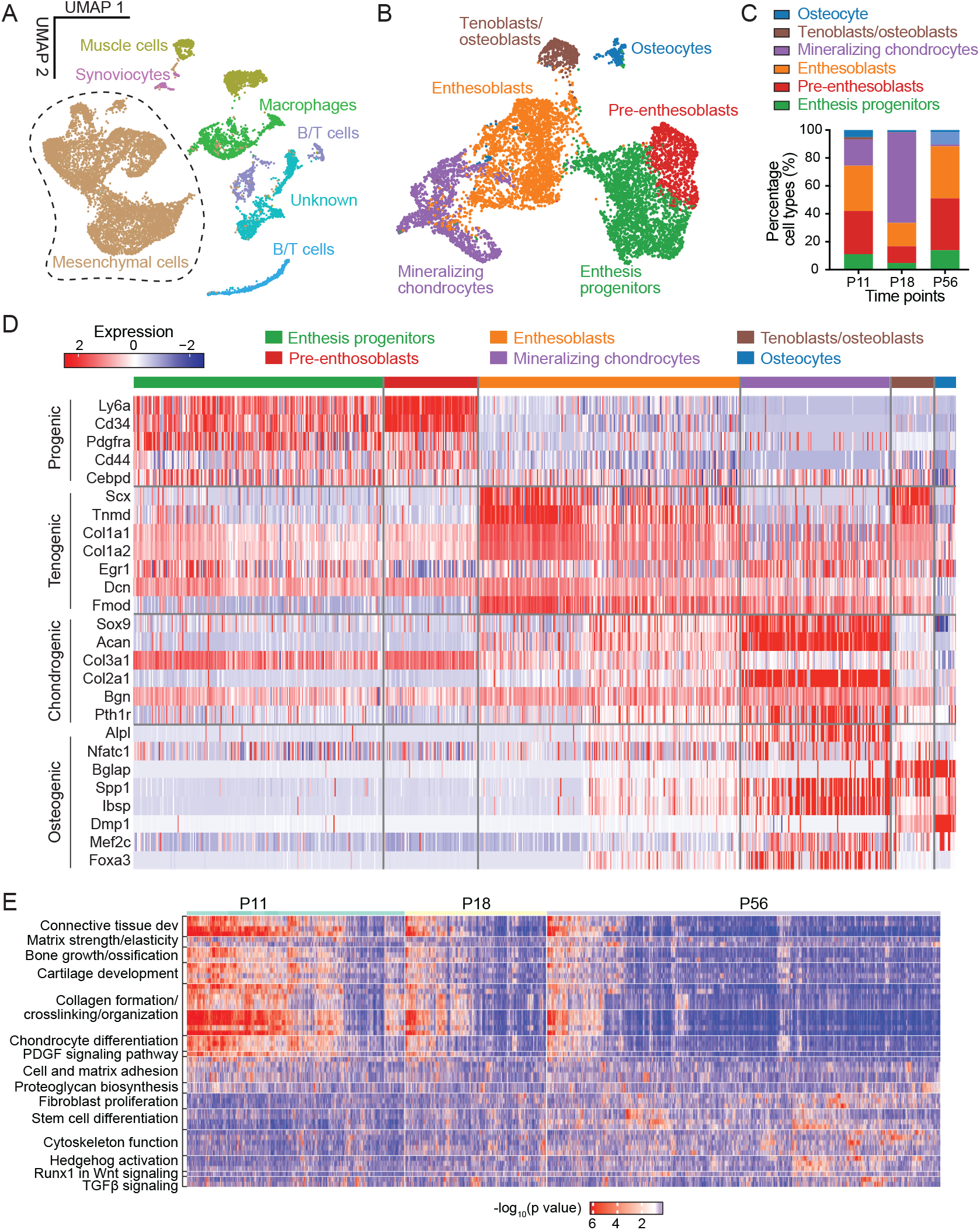
Categorization and transcriptomes of supraspinatus tendon enthesis cells by scRNA- Seq. (A) UMAP plot of all cells integrated from tendon entheses at postnatal day 11 (P11), P18, and P56. (B) Subsetting and further clustering of enthesis mesenchymal cells indicated by dashed outline in (A). Only this subset of enthesis cells were used for the subsequent analyses. (C) Percentage of cell types for P11, P18, and P56. (D) Heatmap of enthesis mesenchymal cells, integrated from P11, P18, and P56, with columns representing each cell subpopulation and rows representing established maker genes for the particular cell types indicated on the left. (E) Heatmap of biological processes at different time points.

Enthesis progenitors had relatively high expression of Ly6a, Cd34, Cd44, and Pdgfra, in agreement with fibro-adipogenic progenitors previously found in tendon, bone marrow, muscle, and colon (De Micheli et al., 2020; Degirmenci et al., 2018; Harvey et al., 2019; Tikhonova et al., 2019), (Figures 1D and S2B). Pre-enthesoblasts were identified and defined based on their clear stem cell transcriptional signatures, but at depressed levels relative to enthesis progenitors, particularly for stem cell maintenance and activated differentiation and transition (Figure S2C). A unique cell subpopulation co-expressing tenogenic (e.g., Scx, Col1a1) and chondrogenic markers (e.g., Sox9, Acan), consistent with previous reports of late fetal enthesis cells (Blitz et al., 2013), were defined as enthesoblasts. We also identified signature profiles of mineralizing chondrocytes (e.g., Sox9, Acan, Col2a1, Alpl, Spp1, Ibsp), tenoblasts/osteoblasts (e.g., Scx, Tnmd, Col1a1, Nfatc1, Bglap), and osteocytes (e.g., Nfatc1, Bglap, Spp1, Dmp1). All cell subpopulations were actively expressing their corresponding signature genes and deactivating other listed gene sets at P18 (Figure S2B). To determine cellular functions across varied cell subpopulations or development stages, we then performed single-cell gene set enrichment (ssGSEA) analysis. The biological processes matrix strength, bone growth, cartilage development, collagen formation, and chondrocyte differentiation were significantly more enriched at P11 and P18 compared to P56 (Figure 1E). The biological processes fibroblasts proliferation, cytoskeleton function, stem cell differentiation, and Runx1 in Wnt signaling were significantly more enriched at P56 compared to the earlier timepoints. Enthesoblasts had the most enriched profiles of matrix deposition and tissue development. These findings demonstrate six distinct enthesis cell populations at early postnatal timepoints that activate several biological processes to build and mineralize the enthesis.

### Candidate regulators for enthesis progenitor cell differentiation and mineralization defined by single-cell network inference (SCENIC) and fluorescence in situ hybridization (FISH)

To determine the gene regulatory network that defines enthesis maturity and cell fate, we applied SCENIC analysis to the enthesis mesenchymal cell data (Aibar et al., 2017). This analysis revealed that P11 and P18 enthesis cells had enriched transcription factor regulons related to tissue mineralization compared to P56 enthesis cells (i.e., Sox9, Runx2, Mef2a, Mef3c, Foxa3, and Nfatc1 regulons; Figure 2A). Consistent with this finding, expression of the master transcription factors for these regulons, such as Sox9, Mef2a, Mef2c, and Foxa3, were upregulated in mineralizing chondrocytes, particularly compared to tenoblasts/osteoblasts and osteocytes (Figure S3), implying that the mineralizing chondrocyte was the major cell subpopulation responsible for enthesis mineralization. Interestingly, Cebpd and Myc regulons were upregulated at P56 compared to the earlier timepoints, implying their possible roles in driving enthesis maturity. The regulons and associated master transcription factors specific for stem cell maintenance and differentiation (i.e., Creb5, Bcl3, Egr1, Tcf4, and Creb3l1) were generally enriched for enthesis progenitors, pre-enthesoblasts, and enthesoblasts (Figure 2A,B, S3). The Creb3l1 regulon was enriched only in enthesoblasts, revealing a putative factor unique to enthesis development. This Creb3l1 regulon includes potential tenogenic target genes (e.g., Scx, Mkx, Tnc, Col1a1, Fmod) as binding sites for Creb3l1 (Figure 2B). Finally, SCENIC analysis also detected enriched Sox9 and Runx2 regulons underlying enthesis development and mineralization, in line with previous reports demonstrating essential roles for Sox9 and Runx2 in cartilage and bone development, respectively (Long and Ornitz, 2013). The identification of the master regulators driving enthesis mineralization and maturity provides a rich set of hypothesis-generating results for future study. To corroborate the scRNA-seq findings and evaluate spatiotemporal expression levels of key representative genes in the enthesis, we performed single molecule fluorescence in situ hybridization (FISH) to examine mRNA of progenic markers (i.e., Cebpd, Ly6a, Cd34), and osteogenic markers (i.e., Nfatc1, Mef2a, Runx2, Foxa3). Tendon enthesis sections were analyzed at P7, P18, and P56. Consistent with scRNA-seq results (Figure 2C), FISH demonstrated positive expression of these genes and revealed their spatial expression patterns (Figures 2D and S4). Cebpd expression was greater in the enthesis at P18 than at P7 and P56. In contrast, Ly6a expression showed low levels of expression across all three timepoints, whereas Cd34 expression increased with time. This contrasts with scRNA-seq analysis, which showed similar expression levels for Cd34, Ly6a, and Cebpd in enthesis progenitors and pre-enthesoblasts across timepoints. This may be due to the different approaches and normalization schemes; for FISH analysis, cell numbers positive for expression were normalized by the total number of enthesis cells, whereas scRNA-seq analysis compared gene expression *levels* for each cell subpopulation. Additionally, expression patterns of these progenic markers were not dependent on enthesis region. For osteogenic markers, Nfatc1 expression was consistent across all timepoints, whereas Foxa3 expression increased with enthesis maturity. Of note, Mef2a and Runx2 expressions were more abundant at P18 than P7 and P56, consistent with the greatest number of mineralizing chondrocytes at P18 (Figure 1C). As expected, osteogenic markers, such as Mef2a Runx2, and Foxa3, were more highly expressed at the zones of mineralized fibrocartilage and bone than tendon.

**Figure 2.**
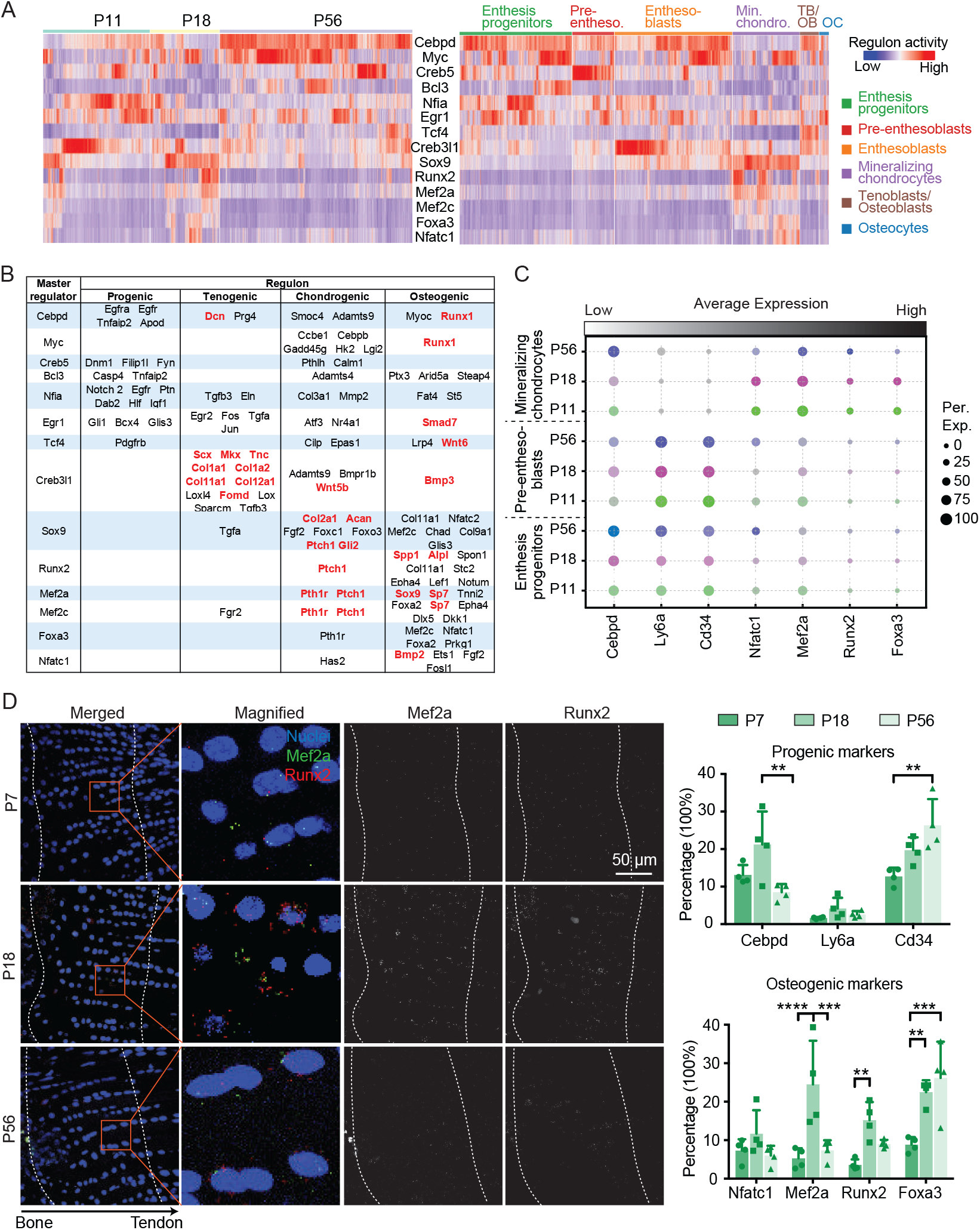
Identification of candidate regulators for enthesis development and mineralization using single-cell network inference and fluorescence *in situ* hybridization (FISH) (A) The activities of select top regulons (rows) presented by time point (left) and cell type (right), as detected by SCENIC analysis. Pre-entheso., pre-enthesoblasts; Min. chondro., mineralizing chondrocytes; TB/OB, tenoblasts/osteoblasts; OC, osteocytes. Pre-entheso., pre-enthesoblasts; Min. chondro., mineralizing chondrocytes; TB/OB, tenoblasts/osteoblasts ; OC, osteocytes. (B) Summary of putative transcription factor regulons (rows) mediating progenitor function, tenogenesis, and chondrogenesis. (C) Average expression levels of transcription factors identified in in (A) and (B) for enthesis progenitors, pre-enthesoblasts, and mineralizing chondrocytes at P11, P18, and P56. The color represents different time points; the brightness of each dot represents the average expression level from low (light) to high (dark); and the size of each dot represents the percentage of positive cells for each gene. (D) Representative FISH images (left) of transcription factors identified at different time points and semi-quantitative histomorphometric analysis of expression of these transcription factors (percentage of cells with positive staining; right). The panels in the second column (left) are magnified images corresponding to the red rectangles. The region between two dash lines denotes the enthesis; *p<0.05, **p<0.01, ***p<0.001.

### Enthesis stem cell differentiation trajectories reveal a path from enthesis progenitors to mineralizing chondrocytes

To explore the origin and lineage trajectories of enthesis mesenchymal cells, we applied Monocle 3 (Trapnell et al., 2014) and ScVelo (Bergen et al., 2020) algorithms to the scRNA-seq data to infer differentiation paths. Monocle 3 uses an unsupervised algorithm, without prior knowledge, to order whole-transcriptome profiles of single cells. The RNA velocity algorithm (Trapnell et al., 2014), on the other hand, predicts future cell states based on the time derivative of the gene expression pattern. Cross-referred results from the two algorithms, therefore, provides a robust approach for determining differentiation trajectories. Partition-based graph abstraction (PAGA) maps from ScVelo analysis suggested that both pre-enthesoblasts and enthesoblasts originated from enthesis progenitors, and enthesoblasts had a greater chance to branch into mineralizing chondrocytes and tenoblasts/osteoblasts than osteocytes (Figure 3A). Strikingly, projection of all cells along the latent time of RNA velocity revealed a clear separation of P56 enthesis cells from P11 and P18 enthesis cells (Figure 3B and S5A). This demonstrates a strong connection between P11 and P18 enthesis cell RNA velocities, whereas P56 enthesis cells may have reached homeostasis and their expression profiles are distinct from the earlier postnatal stages. Ly6a and Cd34, considered progenitor markers, had relatively higher expression levels and RNA velocities in enthesis progenitors (Figure 3C and S5B). Similarly, tenogenic markers (i.e., Scx, Tnmd), chondrogenic makers (i.e., Sox9, Col X), and osteogenic markers (i.e., Runx2, Ibsp, Foxa3) were enriched in corresponding cell subpopulations with respect to gene expression levels and RNA velocities (Figure 3C, S5B and S5C). Enthesoblasts, tenoblasts/osteoblasts, mineralizing chondrocytes, and osteocytes were selected to estimate trajectories of enthesis mesenchymal cell differentiation using RNA velocity (Figure 3D). Mineralizing chondrocytes, tenoblasts/osteoblasts, and osteocytes were rooted in enthesoblasts. Consistent with this ScVelo analysis, Monocle analysis also inferred that enthesis progenitors were the original cell population leading to pre-enthesoblasts and enthesoblasts, which then branched into either mineralizing chondrocytes, tenoblasts/osteoblasts, and enthesoblasts (Figure 3E). Integration of data from all three timepoints and ordered along pseudotime by Monocle revealed a clear temporal trajectory from enthesis progenitors to pre-enthesoblasts, then enthesoblasts, and finally mineralizing chondrocytes (Figure 3E). Overall, comparison of trajectory analyses from Monocle and RNA velocity revealed clear lineage trajectories of enthesis mesenchymal cells originating from enthesis progenitors.

**Figure 3.**
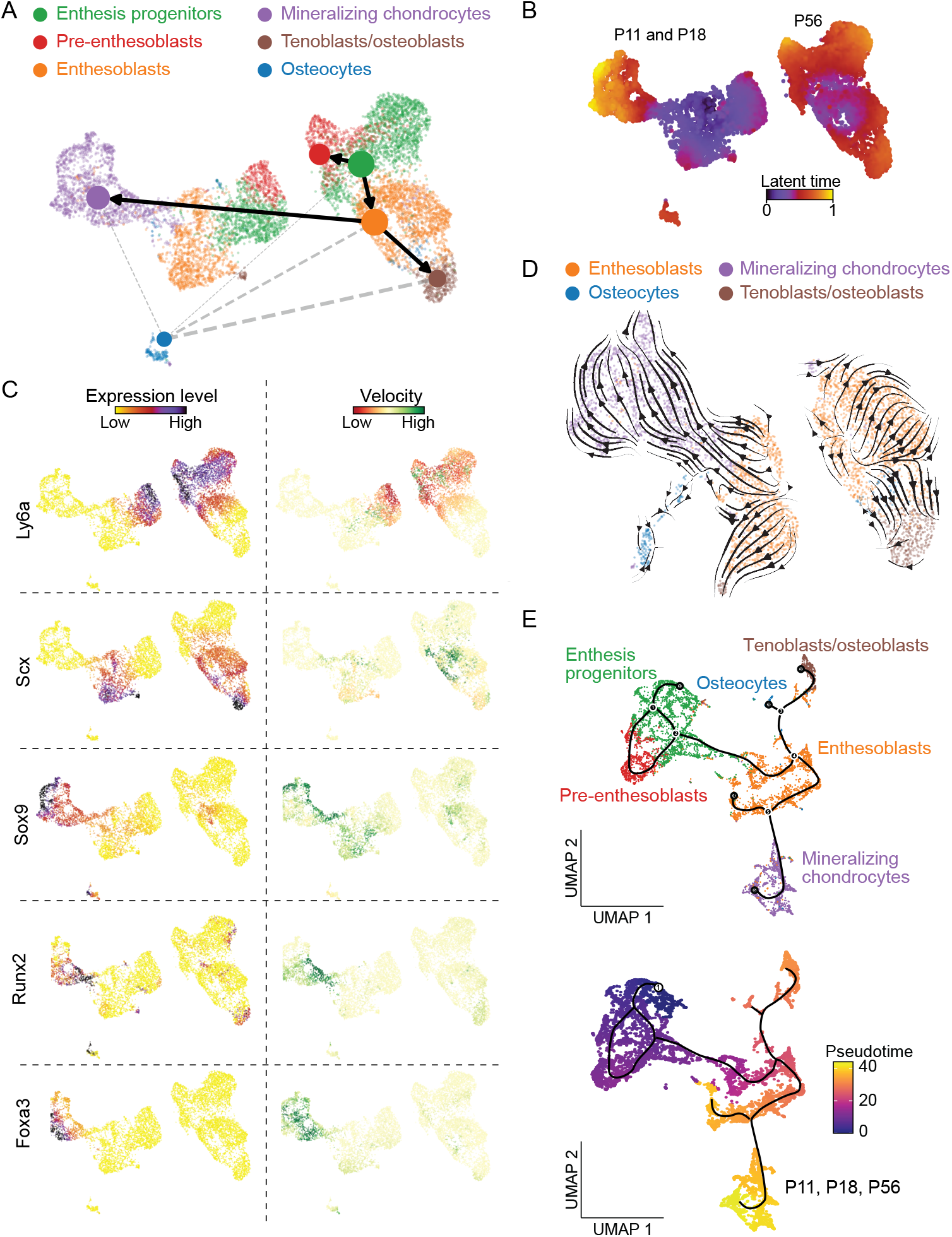
Cell differentiation trajectories of tendon enthesis cells predicted by scVelo and Monocle. (A) Partition-based graph abstraction (PAGA) trajectory model of scVelo predicted cell-type transitions with single-cell embedding, colored by annotated cell types. Thicker and solid lines indicate stronger correlations between cell types; the size of nodes indicates the percentage of each cell type. (B) Latent time plot from scVelo analysis captured the temporal dynamics of enthesis transcriptional profiles at different time points. (C) Expression levels and velocities of representative gene markers, evaluated by scVelo. (D) RNA velocity stream of a subset of enthesis differentiated cells (i.e., enthesoblasts, mineralizing chondrocytes, tenoblasts/osteoblasts, and osteocytes) overlaid with UMAP embedding. Each arrow indicates the direction and speed of cell movement. (E) Differentiation trajectories of enthesis mesenchymal cells ordered along pseudotime using Monocle, colored by cell type (top) and pseudotime (bottom).

### Enthesis progenitors demonstrate clonogenicity and multipotency

In our previous work, we described a unique subpopulation of Gli1+ cells that originated *in utero* were distinct from tendon fibroblasts and epiphyseal chondrocytes (Schwartz et al., 2015). These cells were responsible for building and mineralizing the enthesis and were maintained in the mature enthesis. Furthermore, we found that Gli1+ cells were actively involved in healing neonatal enthesis injuries (Schwartz et al., 2017). Motivated by these prior studies and the scRNA-seq analysis of enthesis mesenchymal cells identifying enthesis progenitors, we explored the nature of enthesis-specific Gli1+ cells as a putative resident source of enthesis progenitors. Gli1+ cells were isolated using fluorescence activated cell sorting (FACS) and analyzed using scRNA-seq and *in vitro* clonogenicity and multipotency assays. Integration of scRNA-seq data of Gli1+ cells from seven timepoints spanning the range from neonatal enthesis development to maturity revealed that Gli1+ cells had abundant expression of progenitor markers, including Ly6a, Cd44, Cd34, and Pdgfra (Figure 4A). Clonogenicity was then compared among Gli1+ cells, Gli+Ly6a+Cd44+ cells (Gl1+ cells further sorted by FACS for these two progenitor markers; Figure 4B), and tendon ScxGFP cells (i.e., mature tenocytes, considered a negative control). Gli1+ cells had better capacity to form colonies compared to ScxGFP cells (Figure 4B). Remarkably, Gli1+ progenitors with expression of Ly6a and Cd44 were seven times as clonogenic as ScxGFP cells and twice as clonogenic as Gli1+ cells. To evaluate whether Gli1+ cells were the source of mesenchymal cells at the enthesis, scRNA-seq data of isolated Gli1+ cells were integrated with data from all enthesis mesenchymal cells. Gli1+ cells were distributed across the entire UMAP of all enthesis mesenchymal cells, suggesting that Gli1+ cells have the capacity to differentiate into all enthesis cell phenotypes (Figure 4C). To examine multipotency, Gli1+ cells, bone marrow-derived mesenchymal stem cells (MSCs, positive control), and tendon ScxGFP cells (negative control) were cultured under adipogenic, osteogenic, and chondrogenic conditions (Figure 4D). Like MSCs, Gli1+ cells had both osteogenic and chondrogenic potentials. However, they did not show adipogenic capacity. This suggests that enthesis Gli1+ cells have primed and irreversible differentiation potential, possibly defined by their specific niches within the enthesis (Harvey et al., 2019). Importantly, differentiated chondrocyte pellets from Gli1+ cells but not MSCs and ScxGFP cells, had the morphology, lacunae, and deposited surrounding matrix similar to that of tendon enthesis, suggesting that Gli1+ cells could serve as a source for enhanced enthesis healing.

**Figure 4.**
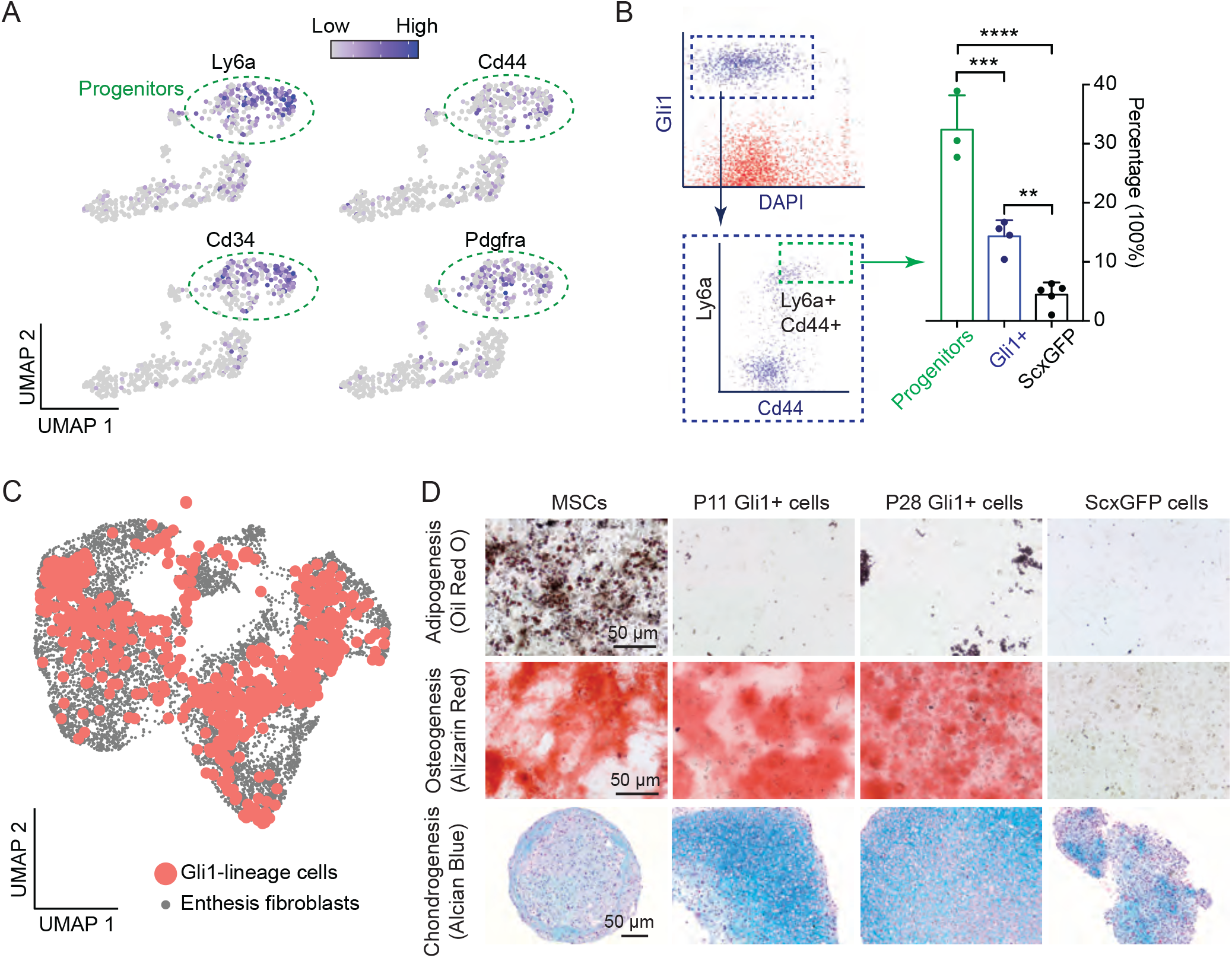
Enthesis progenitors demonstrate clonogenicity and multipotency. (A) UMAP plots of expression levels of progenitor makers Ly6a, Cd34, Cd44, and Pdgfra for enthesis Gli1-lineage (Gli1+) progenitor cells. (B) Gli1+ progenitors, labeled for progenitor markers Ly6a and Cd44 and subjected to FACS had the highest clonogenicity. *p<0.05, **p<0.01, ***p<0.001, ****p<0.0001. (C) UMAP plot of Gli1+ cells from all experiments (Gli-lineage cells from P11, P14, P18, P21, P28, and P42, and enthesis mesenchymal cells from P11, P18, and P56). (D) Enthesis Gli1+ cells showed the capacity for osteogenesis and chondrogenesis, but not for adipogenesis.

### Enthesis Gli1-lineage cells have a linear differentiation trajectory

A lineage-tracing approach was taken to characterize the path of enthesis cells from progenitor through mature phenotypes. Gli1+ enthesis cells were tracked by injecting Gli1Cre^ERT2^;Ai14 mice with tamoxifen at P5 and P7 (Figure 5A). Our prior work demonstrated that this approach is effective in identifying enthesis lineage cells through skeletally mature stages (Schwartz et al., 2017; Schwartz et al., 2015). Gli1+ cells were isolated from microdissected entheses and sorted by FACS. After quality control and filtering, as described in the methods section, Gli1+ cells from seven timepoints were integrated and grouped into five clusters (Figure 5B). Gli1+cells had a significant cluster of enthesis progenitors marked by Ly6a and C34, concordantly with the previous analysis of all enthesis cells (Figures 1D and 2C). The composition of the cell clusters did not change over time, indicating that descendants of Gli1+ cells with different phenotypes were evenly distributed across the enthesis (Figure 5C and S6A). As expected, each Gli1+ cell cluster expressed the expected markers: progenic (i.e., Ly6a, Cd34), tenogenic (i.e., Scx, Tnmd), chondrogenic (i.e., Sox9, Acan), and osteogenic (i.e., Nfatc1, Lbsp) (Figure 5D). The pseudotime trajectory of Gli1+ cells by Monocle 3 analysis revealed a linear lineage progression from enthesis progenitors to pre-enthesoblasts, enthesoblasts, chondrocytes, and finally to mineralizing chondrocytes (Figure 5E). The trajectory analysis was further corroborated by enriched expression of Ly6a only in enthesis progenitors, Col1a1 and Fmod only in enthesoblasts, and Col2a1 and Sox9 only in chondrocytes and mineralizing chondrocytes (Figure 5F).

**Figure 5.**
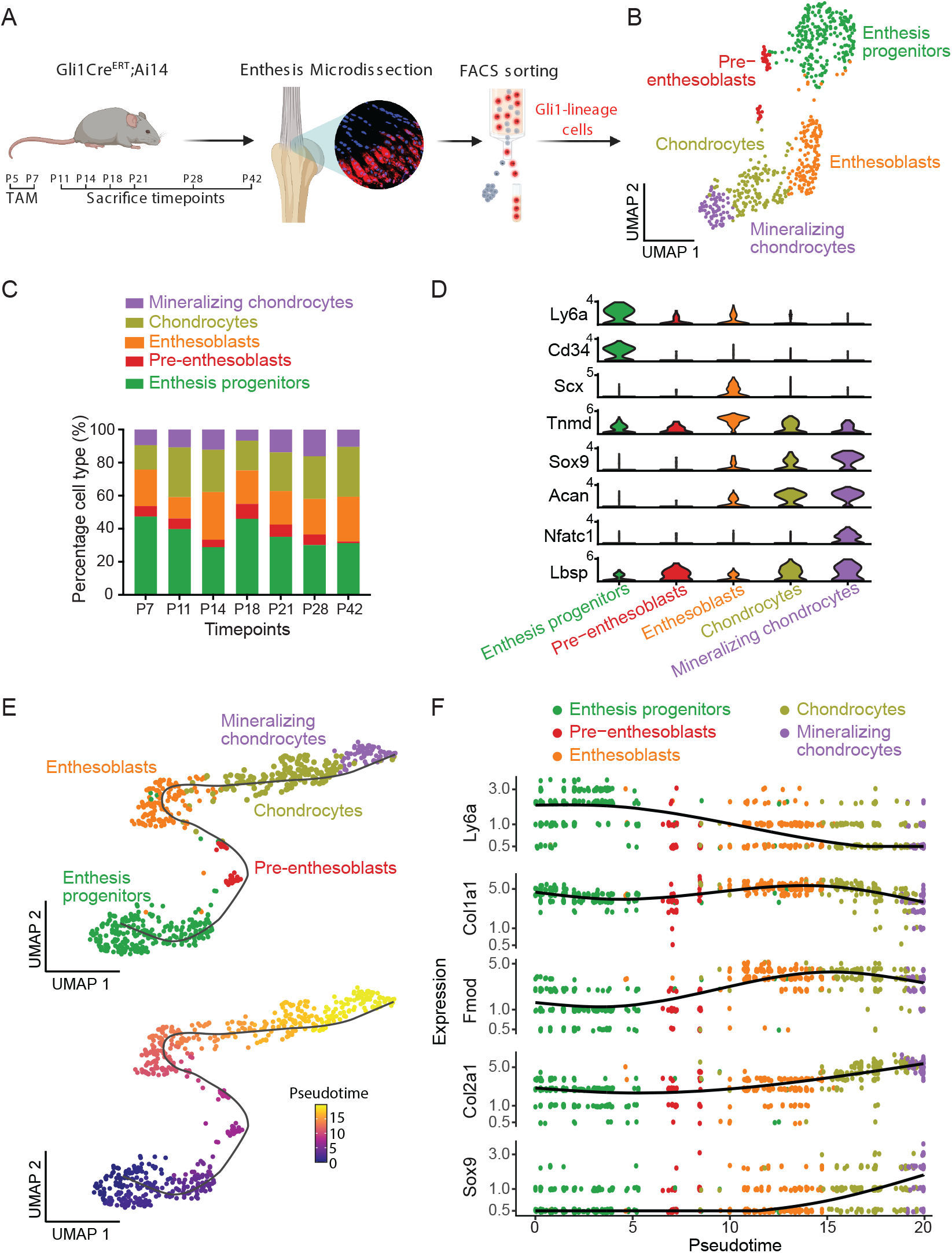
Enthesis Gli1+ cells demonstrate a linear differentiation trajectory. (A) Experimental design for scRNA-seq analysis of enthesis Gli1+ cells from P7, P11, P14, P18, P21, P28, and P42. (B) UMAP plot of Gli1+ cells from GliCre^ERT^;Ai14 mice colored by cell type. (C) Composition of each cell cluster of Gli1+ cells across different time points. (D) Violin plots of gene expression of established markers for annotating cell clusters. (E) A linear trajectory of enthesis Gli1+ cells along pseudotime was identified by Monocle, colored by cell type (top) and pseudotime (bottom). (F) Scatterplots of expression levels of representative genes ordered along pseudotime.

### Chondrocytes and mineralizing chondrocytes are more involved in enthesis formation and maturity

To more deeply evaluate the biological processes driving Gli1+ cell differentiation, single sample gene set enrichment (ssGSEA) analysis was performed (Chen et al., 2019). Among the five clusters of Gli1+ cells, chondrocytes and mineralizing chondrocytes showed the highest activity, particularly for processes related to bone growth and ossification, cartilage development, collagen formation, and chondrocyte differentiation (Figure 6A). This suggests that chondrocytes and mineralizing chondrocytes are the key contributors to extracellular matrix deposition and mineralization. We then performed SCENIC analysis to this dataset to explore transcription factor regulons responsible for mediating Gli1+ cell differentiation and focused on transcription factor regulons related with phenotype shifts and maintenance of enthesis progenitors and mineralizing chondrocytes (Figure 6B). Cebpd expression and regulons were more enriched in enthesis progenitors than in the other cell clusters (Figures 6B and 6C). Jun and Egr1 regulons were also more activated in enthesis progenitors. Distinct from the Egr1 regulon, downstream genes of Cebpd regulons were related to progenitor function and tenogenesis (Figure S6C). Although Pou3f3 had relatively low gene expression, the Pou3f3 regulon was highly activated in chondrocytes and mineralizing chondrocytes (Figures 6B and C). As shown by the scRNA-seq analysis of all enthesis cells, both expression and regulon activities of Nfatc1 and Foxa3 were enriched in chondrocytes and mineralizing chondrocytes. Expression patterns of progenic and osteogenic markers were confirmed across development stages (Figure 6D). Progenic markers, such as C34, Ly6a, and Cebpd were down- regulated at P7 compared to late postnatal timepoints in progenitors. This was also the case for osteogenic markers Foxa3, Mef2a, and Nfatc1. Of note, the expression of Mef2a, which has been reported to regulate bone formation by controlling SOST in osteocytes (Leupin et al., 2007), increased with enthesis maturity from P7 to P42.

**Figure 6.**
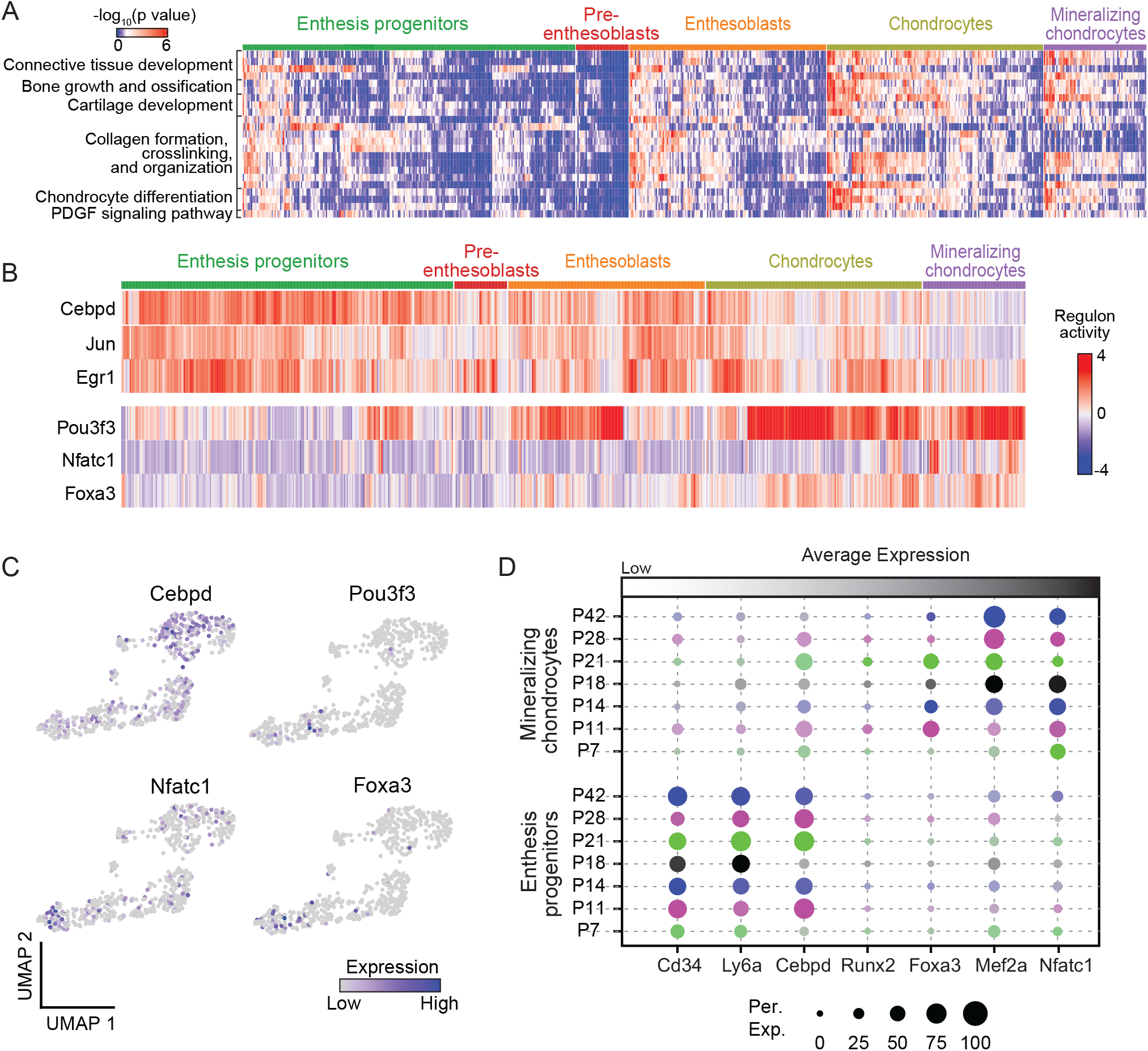
Transcriptional regulation of Gli1+ cells. (A) Heatmap of biological processes of Gli1+ cells grouped by cell clusters. (B) Activities of selected top regulons for Gli1+ cells organized by cell clusters, as identified by SCENIC analysis. (C) UMAP plots of expression levels of identified transcription factors for Gli1+ cells. (D) Average expression levels of representative gene makers for enthesis progenitors and mineralizing chondrocytes of Gli1+ cells at different time points. The color represents different time points; the brightness of each dot represents the average expression level from low (light) to high (dark); the size of each dot represents the percentage of positive cells.

### Therapeutic potential of Gli1+ progenitor cells for enthesis healing

Gli1+ cells are necessary to build the enthesis, express progenitor cell markers, and show multipotency and clonogenicity *in vitro*. We therefore explored the therapeutic potential of Gli1+ cells to enhance enthesis healing, with the long-term goal of developing a stem-cell-based treatment for enthesis regeneration. Gli1+ cells were sorted from mouse entheses, expanded *in vitro,* and transplanted to injured entheses (Schwartz et al., 2017). Gli1+ cells engrafted onto the injury site and were retained from post- operative day 1 (POD1) to POD7 (Figure S7D). Enthesis injury defects became smaller with time, with deposition of new extracellular matrix (Figures 7A). Enthesis injuries treated with Gli1+ cells had smaller gaps and greater mineralization than untreated enthesis injuries. Quantification of microCT data showed that bone density and volume of injured regions increased with time and were significantly greater in Gli1+ treated injuries compared to control, particularly at POD28 (Figure 7). Trabecular bone thickness in Gli1+ cell-treated injuries was significantly less than control at POD7, but similar at the other timepoints. Histologically, collagen fiber aligned improved and cell density decreased with time (Figure 7A,B). More extracellular matrix (including cartilaginous tissue) was deposited at the injured region in the Gli1+ cell-treated group than control, especially at POD28. Semi-quantitative analysis of histology images showed that there was a progressive reduction in histological score towards normal, with significantly better healing in the Gli1+ cell treated group (Figure 7B, S7F). To verify that the positive effects of Gli1+ cell treatment was due to their enthesis progenitor cell nature, we delivered ScxGFP cells extracted from adult tendons (which do not show progenitor characteristics, Figure 4B,D) to the injury site in a separate group of animals. Treatment with ScxGFP tendon cells showed less organized collagen fibers and decreased cartilaginous tissues at POD11 relative to Gli1+ cell-treated injuries (Figure S7E). The results demonstrate the therapeutic potential for Gli1+ stem cell therapy for enhanced enthesis healing.

**Figure 7.**
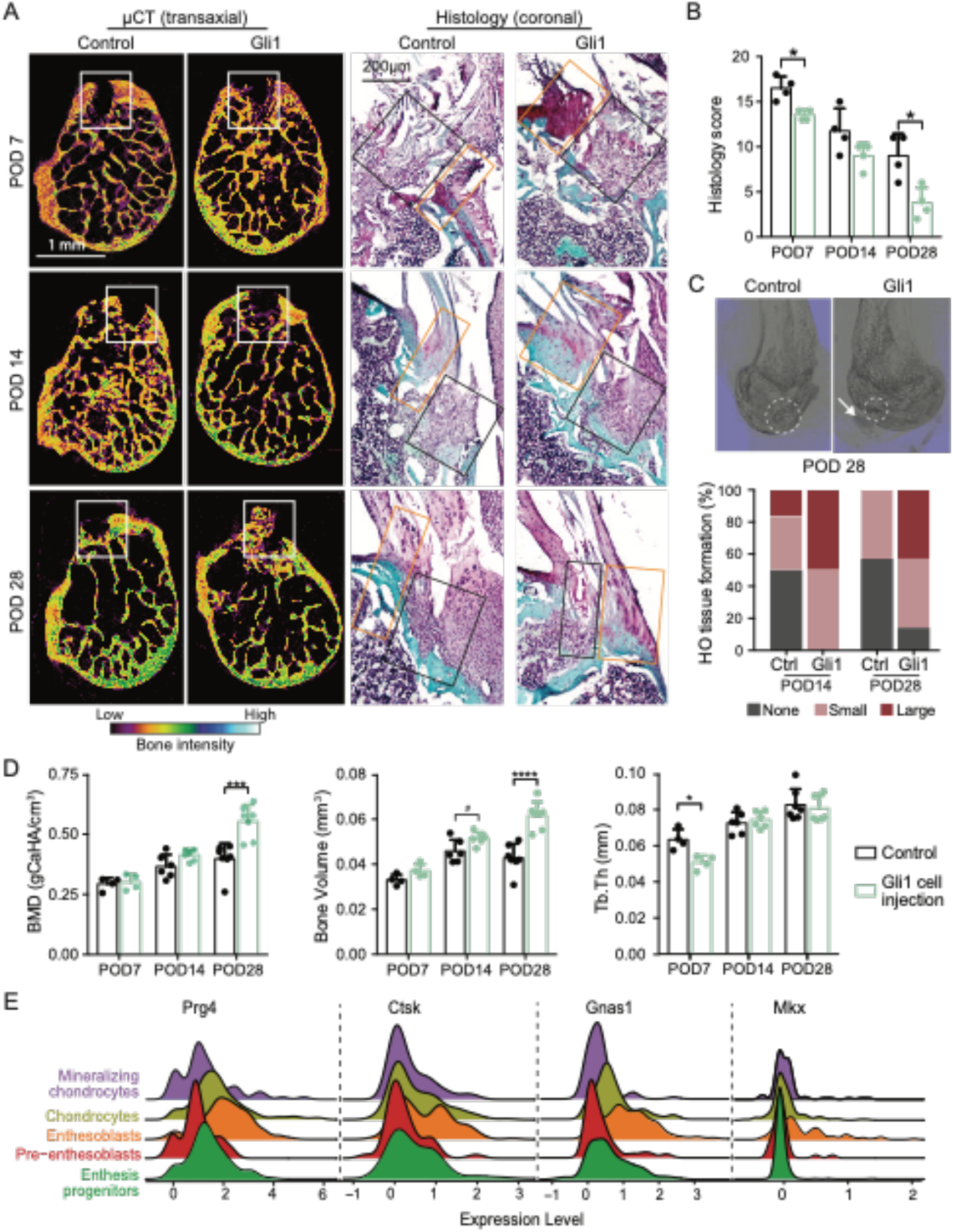
Transplanted Gli1+ cells enhance enthesis healing. (A) Representative μCT sections (left) and histological images (right) of injured entheses at different post- operative days (POD). The white rectangles in the μCT images identify the injured enthesis regions; the black and yellow rectangles in the histological images show injured and intact enthesis regions, respectively; the color bar shows bone density from low to high. The histology sections are stained with safranin O. Control, only collagen gel delivered; Gli1, Gli1+ cells delivered via collagen gel. POD7, post- operative day 7. (B) Histological scoring, as determined from blinded evaluation of histological images. *p<0.05. (C) Representative 3D μCT images of the humeral head bone at POD28 (top), with the injured enthesis outlined with a dashed white circle. The percentage of samples with ectopically calcified tissues near the enthesis, normalized by sample size of each group, is shown on the bottom. The white arrow points a region with heterotopic ossification (HO). “None” denotes no or negligible HO tissue formed close to enthesis; “Small” denotes relatively small HO tissue formed; “Large” denotes relatively large (>20 μm) HO tissue formed. (D) Bone quality of the injured regions, including bone density (BMD), bone volume, and trabecular thickness (Tb.Th), quantified from μCT data. *p<0.05, **p<0.01, ***p<0.001, ****p<0.0001. (E) Density plots of expression levels of HO-related genes for all Gli1+ cells integrated from P7, P11, P14, P18, P21, P28, and P42.

Heterotopic ossification (HO) was also found adjacent to injured entheses in many cases (Figures 7C, S7A). More and larger ossified tissues were identified at Gli1+ cell-treated injuries at POD14 and POD28 compared to the control group. To explore the mechanisms driving heterotopic ossification, we correlated the scRNA-seq data of enthesis Gli1+ cells with a list of known gene markers for HO. Prg4 and Ctsk are associated with osteogenic differentiation at subchondral bone and ligament and tendon (Feng et al., 2020; Novince et al., 2012). Gnas, an osteochondral progenitor gene, is associated with osteogenic differentiation through Wnt signaling (Khan et al., 2018). Mutation of Mkx has been reported to cause aberrant differentiation of tendon progenitors leading to tendon ossification (Liu et al., 2019). In the current study, expression of Prg4, Ctsk, Gnas1, and Mkx were more abundant in enthesoblasts than in the other cell clusters. These results highlight that enthesoblasts can serve as a cell source for tendon heterotopic ossification and are a potential target for inhibiting tendon heterotopic ossification in pathologic settings.

## Discussion

The enthesis serves a critical function in the musculoskeletal system: transferring muscle forces from tendon to bone to achieve joint motion. Prior work has focused on understanding tendon enthesis structure, function, and pathology, but the nature and phenotype shift of enthesis stem cells has remained elusive. Here, we report a detailed transcriptional profile of enthesis mesenchymal cells, including characterization of cell types, exploration of the underlying gene regulatory network, and differentiation path of enthesis progenitor cells to mature enthesis cells. A set of transcription factor regulons, defined by analysis of single-cell transcriptome data and localized to the enthesis by FISH, were identified, specifically for progenitor function, enthesis mineralization, chondrogenesis, tenogenesis, and osteogenesis. Importantly, enthesis progenitors were identified, demonstrating expression of Gli1, Cd34, Cd44, and Ly6a and *in vitro* clonogenicity and multipotency. These newly identified enthesis stem cells demonstrated therapeutic potential for improving enthesis healing.

The development of effective treatments for the tendon enthesis requires a deeper understanding of enthesis cell biology, especially regarding the spatial gradient in cell phenotype from unmineralized to mineralized tissue (Gracey et al., 2020). Fibroblast-like stromal cells, typically denoted as “tenocytes”, have been reported as the main cell type in healthy tendon (Fang and Lake, 2016; Millar et al., 2021). These spindle shaped cells are dispersed along collagen fibers and are responsible for maintaining tendon extracellular matrix (Fang and Lake, 2017; Fang and Lake, 2015). The tendon surface contains an additional resident cell population, αSMA-positive myofibroblasts, which respond to injury and infiltrate the torn region (Best et al., 2020; Wang et al., 2017b). Additionally, various immune cell subtypes, whether resident or recruited, are involved in the initiation and progression of tendon degeneration and injury, but the role of this immune cell subpopulation is controversial for healthy and homeostatic tissues (Gracey et al., 2020). Notably, resident tendon stem/progenitor cells have been identified and isolated based on their *in vitro* behavior (Bi et al., 2007). Although the use of tendon stem cells or stem cell-derived exosomes in animal models is attractive, the lack of determined stem cell markers and knowledge about molecular mechanisms driving their behavior hinders their translational application (Millar et al., 2021). To explore these concepts in the enthesis, we used whole transcriptomic profiles to define six cell subpopulations in mouse supraspinatus tendon according to gene markers related to stem cells, tenogenesis, chondrogenesis, and osteogenesis. Enthesis progenitors shared similar markers with fibro-adipogenic progenitors found in tendon (Harvey et al., 2019), bone marrow (Zhong et al., 2020), and muscle (De Micheli et al., 2020). Different from progenitors, pre-entheoblasts had a transitional expressing profile with up-regulated genes involved in cell differentiation and down-regulated genes attributed to maintenance of progenitor status. An interesting cell subpopulation possessing expression patterns of both tenogenesis and chondrogenesis were labeled as enthesoblasts. This is consistent with previous reports that a distinct cell pool, which expresses both the tenogenic marker Scx and the chondrogenic marker Sox9 (and are not derived from chondrocytes), are found at the enthesis during pre-mineralization fetal timepoints (Blitz et al., 2013; Sugimoto et al., 2013). We suggest that another cell subtype, which we termed “mineralizing chondrocyte”, is responsible for the graded structure of enthesis from unmineralized to mineralized fibrocartilage as it showed activated expression profiles of both chondrogenic and osteogenic markers and clearly played an essential role in enthesis mineralization.

Systematic examination of the gene regulatory network defining enthesis cell phenotypes revealed a group of new transcription factors that regulate lineage progression from progenitor to mature enthesis cell. Further *in vitro* screening and *in vivo* animal models are needed to show whether these factors are necessary and sufficient to drive enthesis progenitor cell differentiation. Nevertheless, a set of new gene markers was identified for the various cell subtypes in the tendon enthesis. These markers will allow for examination of cell phenotype shifts in degenerated or diseased conditions, e.g., towards fibrous, cartilaginous, and/or ossified tissues in tendon and enthesis. Corroborating with the literature that Sox9 is required for chondrogenesis, including at the mineralizing growth plate (Akiyama et al., 2002; Harada and Rodan, 2003; Kronenberg, 2003), Sox9 was confirmed in our data as a downstream target gene contributing to enthesis chondrogenesis and osteogenesis. Similarly, Runx2 is known as the earliest determinant of osteoblast differentiation (Wei et al., 2015), and was also found to be highly activated in enthesis mineralizing chondrocytes and osteoblasts. Our work further highlighted that Cebpd, Nfia, and Egr play a role in mediating downstream genes ranging from progenic to osteogenic markers. Egr1 has been shown to coordinate matrix production and further mediate musculoskeletal tissue formation (Havis and Duprez, 2020; Lejard et al., 2011), but the roles of Cebpd and Nfia in musculoskeletal tissues are not clear. Nfia has been suggested to be expressed in the tendon enthesis and to safeguard stem cell identity in hair follicles (Adam et al., 2020; Kult et al., 2021). A discovery in this study is that six typical tenocyte markers (e.g., Scx, Mkx, Tnc, Col1a1, Col1a2, Fmod) were identified to be regulated by Creb3l1, whose role has not been previously reported for tendon development, with only some evidence suggesting its involvement in collagen metabolic process via TGFβ signaling (Chen et al., 2014; Murakami et al., 2009; Tan et al., 2020). Finally, Runx2, Mef2a, Mef2c, Foxa3, and Nfatc1 were found here to regulate downstream target cofactors required for enthesis chondrogenesis and osteogenesis, consistent with prior literature (Asagiri et al., 2005; Ionescu et al., 2012; Leupin et al., 2007b). Therefore, these newly identified enthesis transcription factors and their related downstream cofactors are temporospatially coordinated to drive progenitor commitment to intended cell fates and then produce a spatially graded mineralized structure at the tendon-bone interface.

Gli1+ cells have been identified in stem cell niches of a wide range of tissues. These cells function as stem/progenitor cells responsible for formation and homeostasis of tissues such as bone, teeth, tendon, colon, kidney, and heart (Degirmenci et al., 2018; Kramann et al., 2016; Kramann et al., 2015; Men et al., 2020; Schneider et al., 2017; Shi et al., 2017). The current work demonstrates that Gli1+ cells are also a specialized pool of resident enthesis progenitor cells. This finding has significant implications for the treatment of enthesis disorders and holds great promise for enhancing tendon-to-bone repair. Enthesis Gli1+ progenitors share similar transcriptional signatures with mesenchymal stem cells, such as enriched Ly6a, Cd34, and Pdgfra, and have the capacity to differentiate into multiple cell types. In the context of enthesis development, Gli1+ progenitors have a linear lineage trajectory from progenitor to mature enthesis cell and populate all enthesis cell subtypes across the transition from tendon to bone. Although the overlap of transcriptional and epigenetic profiles of Gli1+ progenitors from various tissues remains unknown, their resident environment might dictate a limited range of phenotypes specific to the particular tissue. For the enthesis, Gli1+ progenitors were only capable of *in vitro* differentiation into chondrocytes and osteoblasts, but not adipocytes.

Development of therapeutic strategies focused on Gli1+ progenitors may have a wide application to various enthesis injury and degeneration scenarios. For enthesitis, suppression of these cells may mitigate the formation of osteophytes. For tendon-to-bone repair, promotion (or delivery) of these cells may lead to regeneration of a functionally graded enthesis. However, several challenges exist for implementing these strategies. Due to the challenges of harvesting sufficient Gli1+ progenitors from tissues, the decrease in Gli1+ cells with age, and the need to expand cells *in vitro* prior for transplantation (with potential loss of cell phenotype), the therapeutic use of primary Gli1+ progenitor cells is unlikely. It will likely be necessary to use mesenchymal stem cells from another source, or induced pluripotent stem cells, differentiated into enthesis-specific Gli1+ progenitors (Lindvall et al., 2004). Achieving this requires a complete understanding of the gene regulatory network that drives stem cells to become the range of enthesis cell phenotypes. Furthermore, careful design of a cell delivery system is necessary to maintain progenitor cell features after transplantation, engraft cells to the injured region, and promote cell integration and directed differentiation. As presented in our study, Gli1+ cell transplantation can increase matrix deposition and mineralization of injured enthesis but can also cause heterotopic ossification; this was attributed to the limited ability of the collagen gel carrier to retain cells at the injured site. These results also imply that undesired differentiation of Gli1+ progenitors during tendon and kidney calcification and bone marrow fibrosis might be inhibited by preventing proliferation and differentiation of these cells (Kramann et al., 2016; Kramann et al., 2015; Schneider et al., 2017). This can be realized by inactivating molecular signals such as the transcription factors identified in the current study, e.g., by regulating hedgehog signaling components such as Ptch1 and Gli1 or using many available antagonists or agonists (Amakye et al., 2013).

## Limitations of Study

Enthesis cells reside within a dense extracellular matrix, much of it mineralized. Therefore, extraction of cells for FACS and scRNA-seq requires a relatively harsh digestion protocol. The digestion environment and mechanical perturbation by sorting could cause a change in the expression profiles of the cells (van den Brink et al., 2017). However, we demonstrated that the application of chemical digestion and FACS- activated cell sorting resulted in more than 75% viability of cells from the enthesis. Furthermore, cells from all isolation timepoints were dissociated using the same protocol so that comparison of their transcriptomes would be valid. Future work will include spatial transcriptomics to address possible dissociation artifacts and gain spatial information. A second limitation in the study was that delivery of Gli1+ cells to the injured enthesis could lead to heterotopic ossification in some cases. This was likely due to the delivery system used, which did not retain all cells at the repair site post-implantation. To avoid heterotopic ossification in tendon, future experiments will use a biomimetic adhesive material to localize and retain the cells to the injury site.

## Acknowledgements

This work was funded by NIH R01 AR055580. We would like to thank Caisheng Lu and Chingyuan Chen from Flow Cytometry Core at Columbia Center for Translational Immunology for helping with FACS. scRNA-seq experiment was conducted in Single Cell Analysis Core of Columbia Genome Center and Biomedical Informatics Resources at Columbia University, supported by the National Center for Advancing Translational Sciences, through an NIH Grant Number UL1TR001873. Confocal imaging was performed in the Confocal and Specialized Microscopy Shared Resource of the Herbert Irving Comprehensive Cancer Center at Columbia University, supported by an NIH grant P30 CA013696 (National Cancer Institute). The authors thank Siyu He for scientific discussion on RNA velocity analysis.

## Author contributions

F.F. and S.T. designed the study. F.F. conducted experiments. F.F. and Y.X. performed the bioinformatics analyses. F.F., Y. X., K.L., and S.T. interpreted the data. F.F., Y. X., K.L., and S.T. wrote the manuscript. All authors edited and reviewed the manuscript.

---

**Figure S1.**
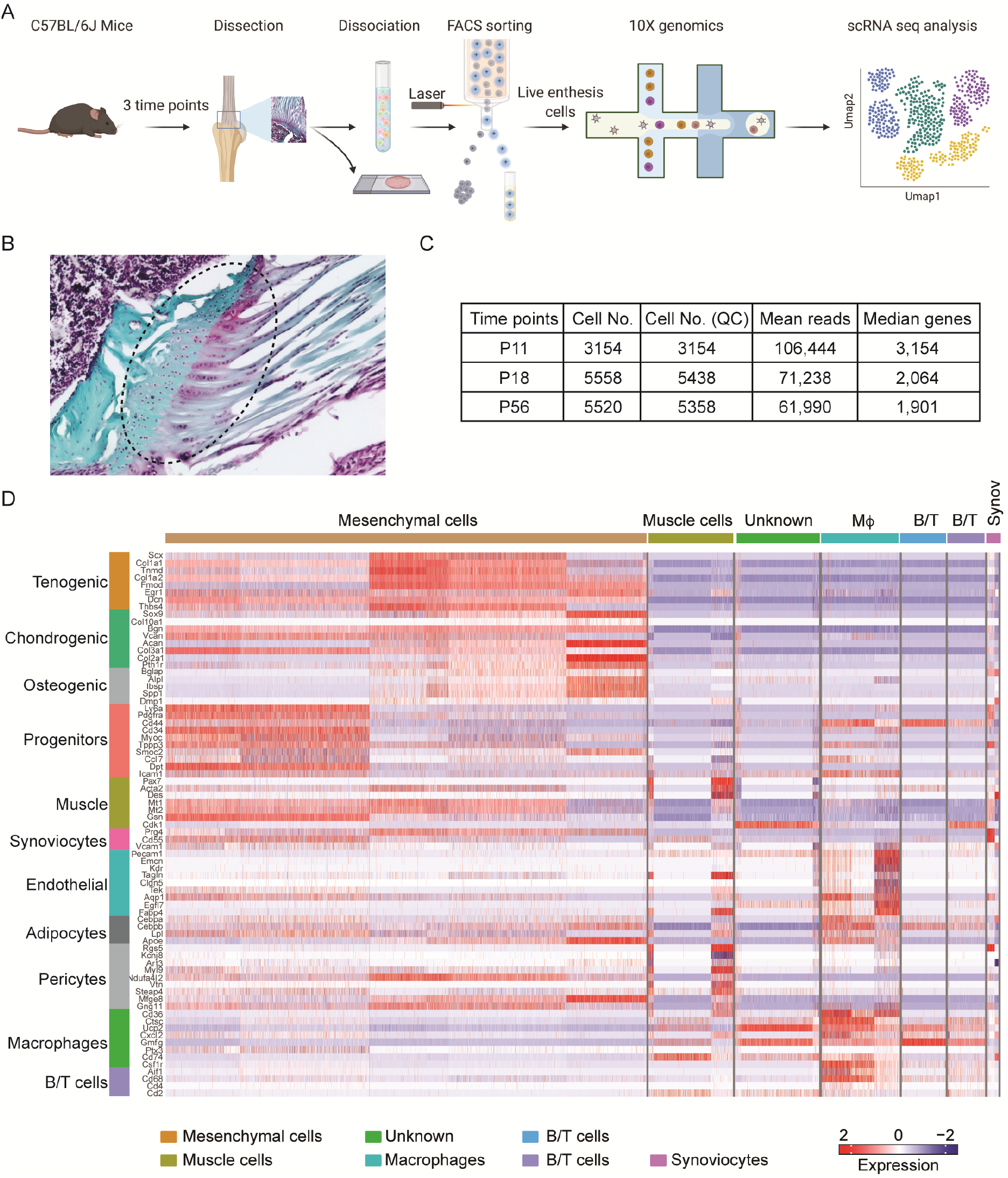
Experimental design and scRNA-Seq analysis of mouse supraspinatus tendon entheses, related to Figure 1. (A) Supraspinatus tendon entheses of C57BL/6J mice at postnatal day 11 (P11), P18, and P56 were pooled, dissociated, and sorted for scRNA-Seq analysis. Enthesis tissues were also evaluated by fluorescent in situ hybridization (FISH) to verify and localize transcription factors of interest. (B) Histological section of a supraspinatus tendon enthesis (stained with Safranin O) showing the region (in dicated by the dashed oval) dissected for scRNA-Seq analysis. (C) Total cell numbers and sequencing reads for scRNA-Seq experiments at each time point. Cell No. (QC) indicates the number of cells that passed quality control. (D) Heatmap showing expression patterns of gene markers for all enthesis cells. This analysis facilitated cell cluster annotation and further selection of mesenchymal cells for subsequent analyses.

**Figure S2.**
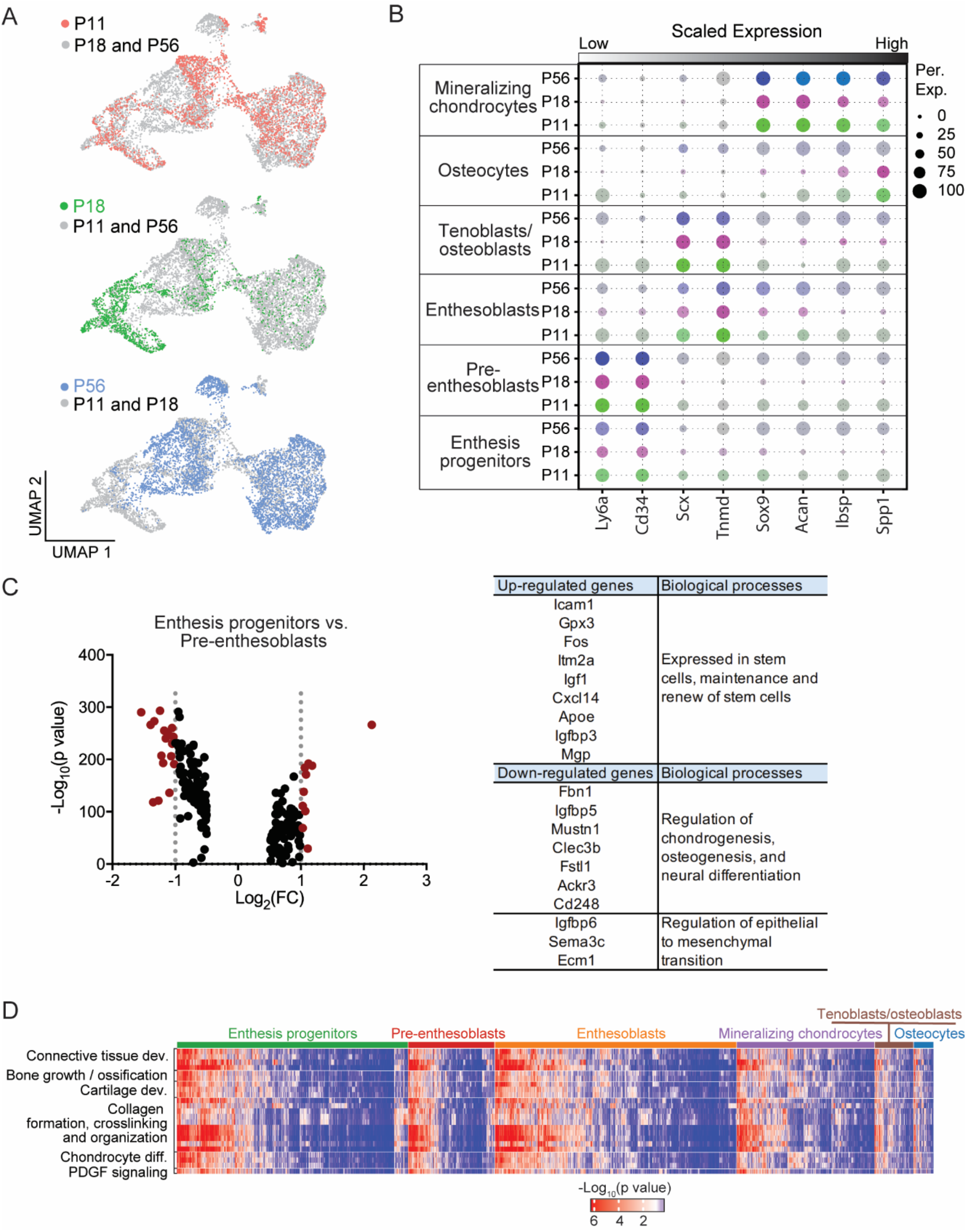
Characterization of enthesis mesenchymal cells for specific gene sets and biological processes, related to Figure 1. (A) UMAP plot displaying cell domains at different time points after integration. (B) Dot plots showing signature gene expression of different cell subpopulations across different time points. The color marks mean gene expression and the dot size indicates the proportion of cells expressing the corresponding gene. (C) Differentiated gene sets of enthesis progenitors, displayed as a volcano plot (left) and table (right), identified by comparing to pre-enthesoblasts. (D) Heatmap revealing the enriched statistical significance of different biological processes for each cell cluster.

**Figure S3.**
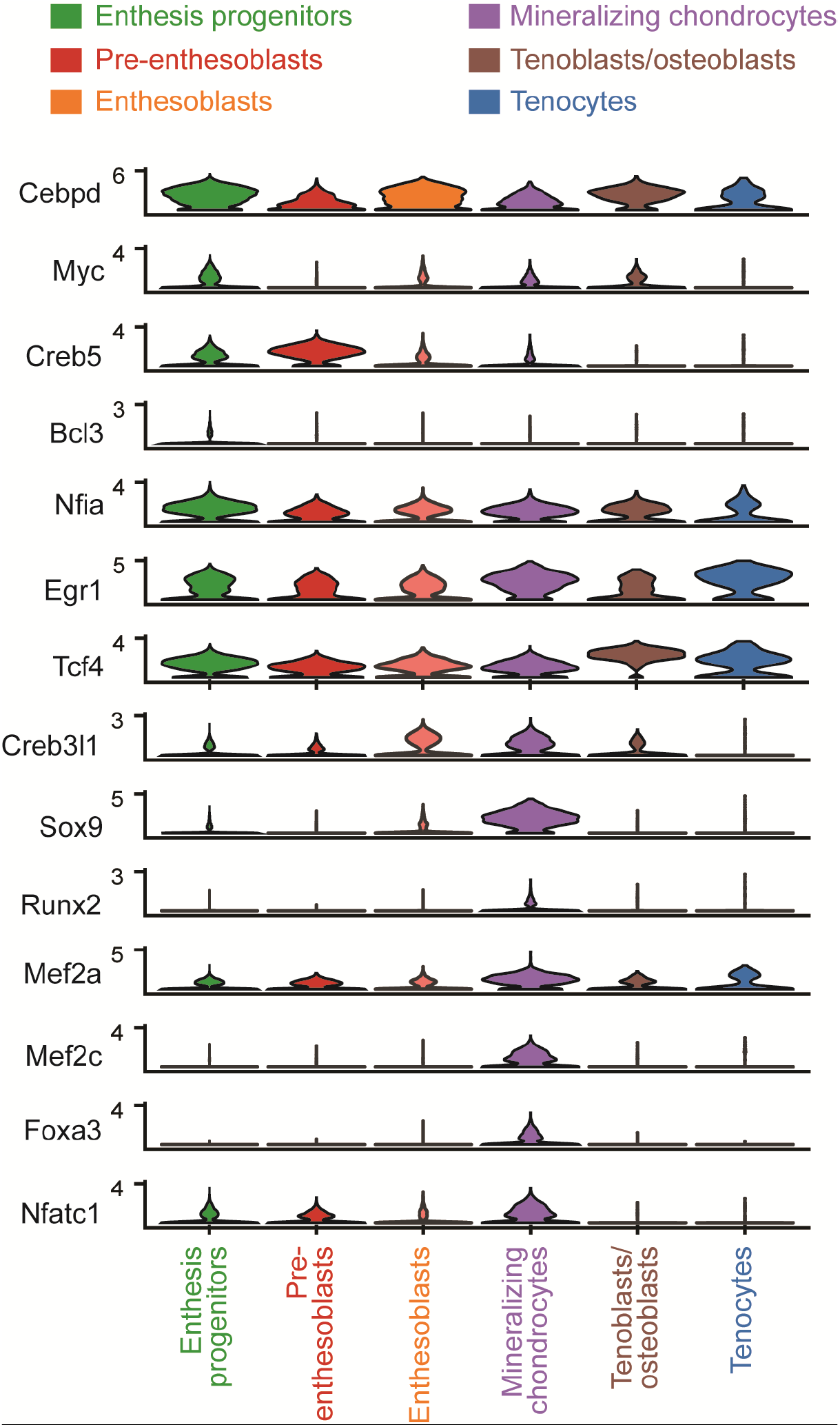
Expression profiles of transcription factors determined by the computational algorithm Scenic, shown as violin plots, related to Figure 2.

**Figure S4.**
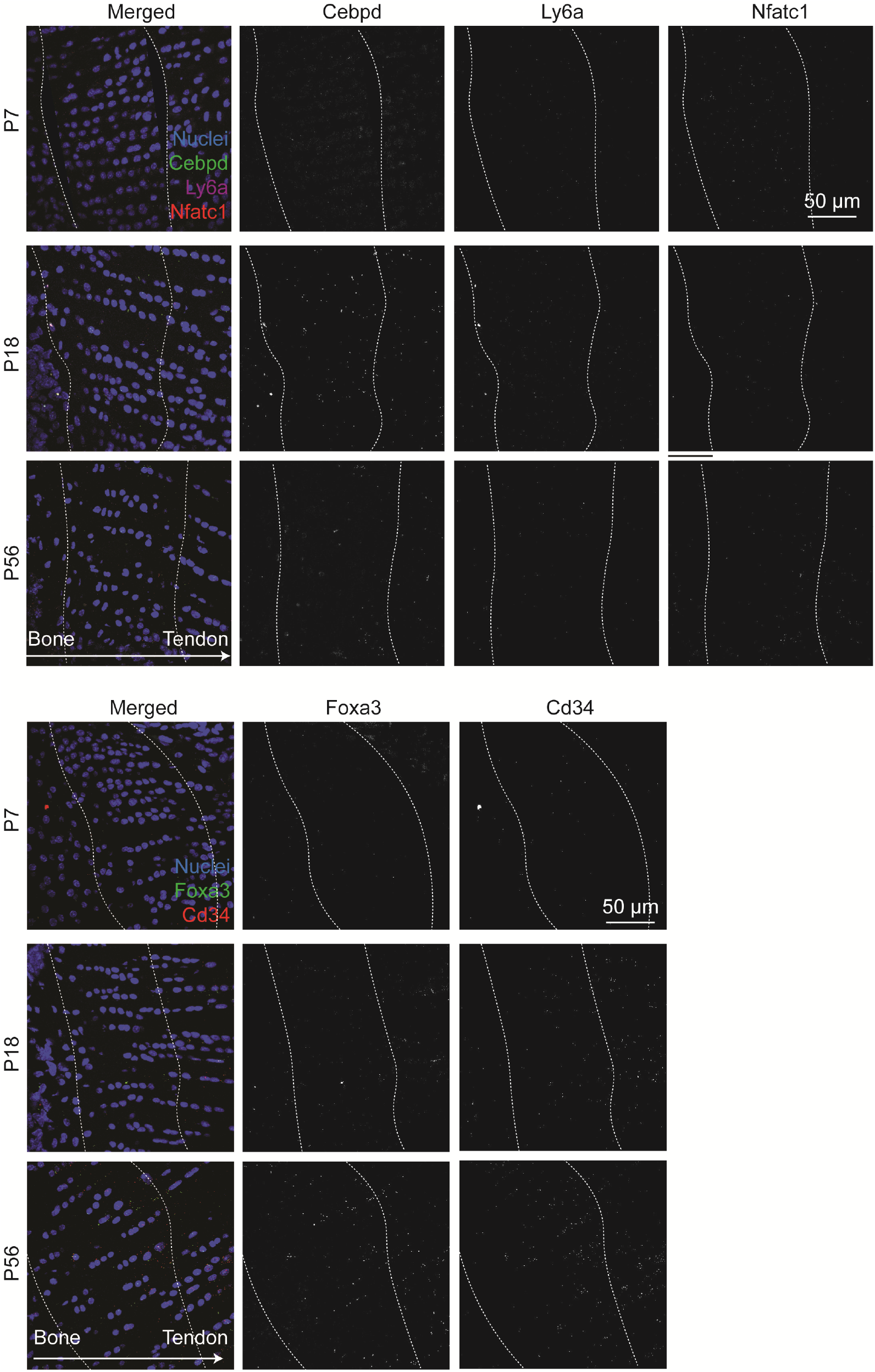
Representative FISH images of supraspinatus tendon entheses showing mRNA expression of transcription factors and gene markers of interest, related to Figure 2. The region between two dash lines denotes the enthesis

**Figure S5.**
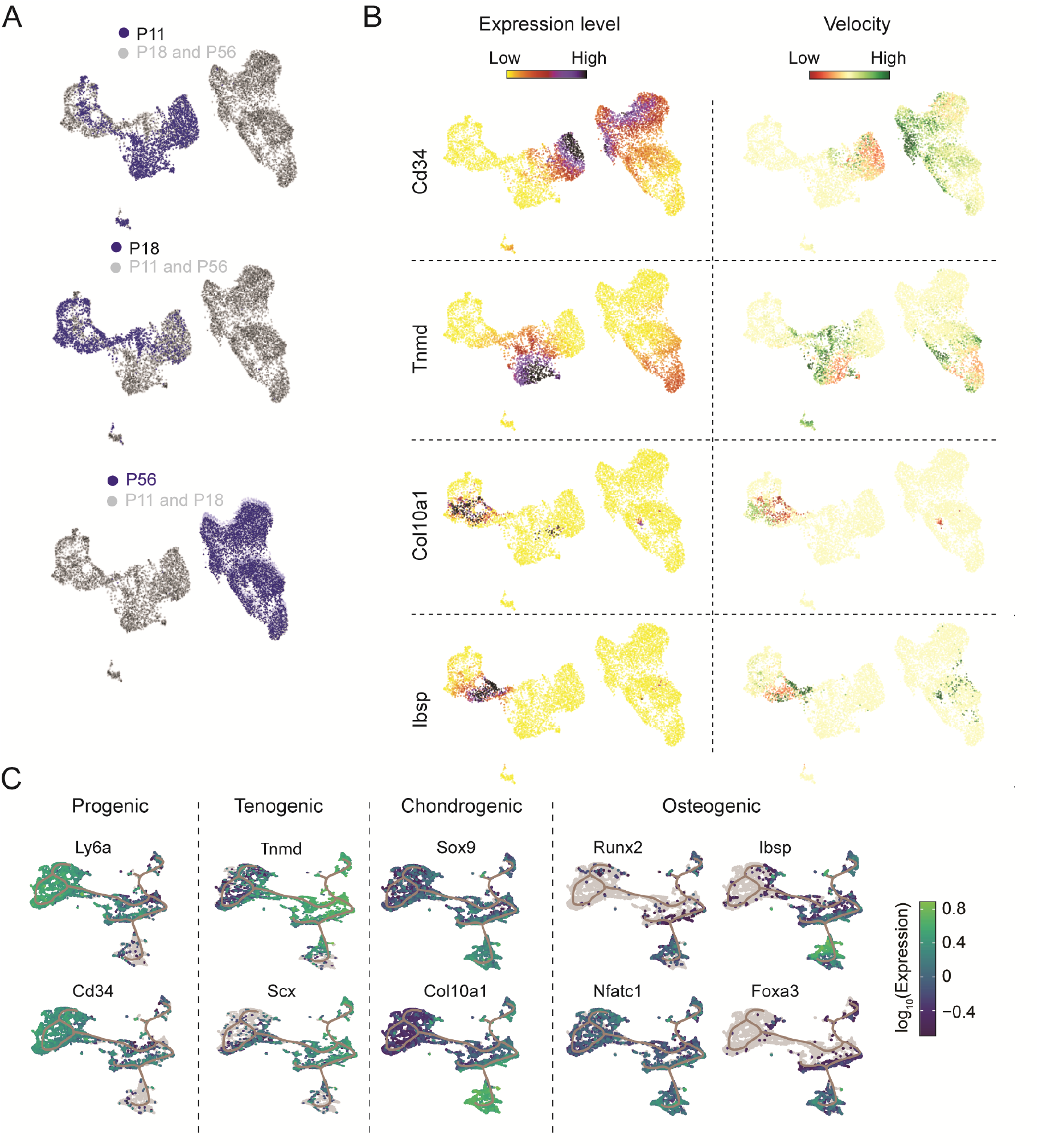
Lineage trajectory analysis of enthesis cells from P11, P18, and P56, related to Figure 3. (A) Enthesis cells from each time point projected to partition-based graph abstraction (PAGA) maps. (B) Expression levels and corresponding velocities of selected genes for enthesis cells integrated from P11, P18, and P56, as determined by scVelo analysis. The color spectrums indicate expression levels and velocities, from low to high. (C) Expression levels of progenic, tenogenic, chondrogenic, and osteogenic gene markers for enthesis cells integrated from P11, P18, and P56, aligned along the trajectories, as determined by Monocle 3.

**Figure S6.**
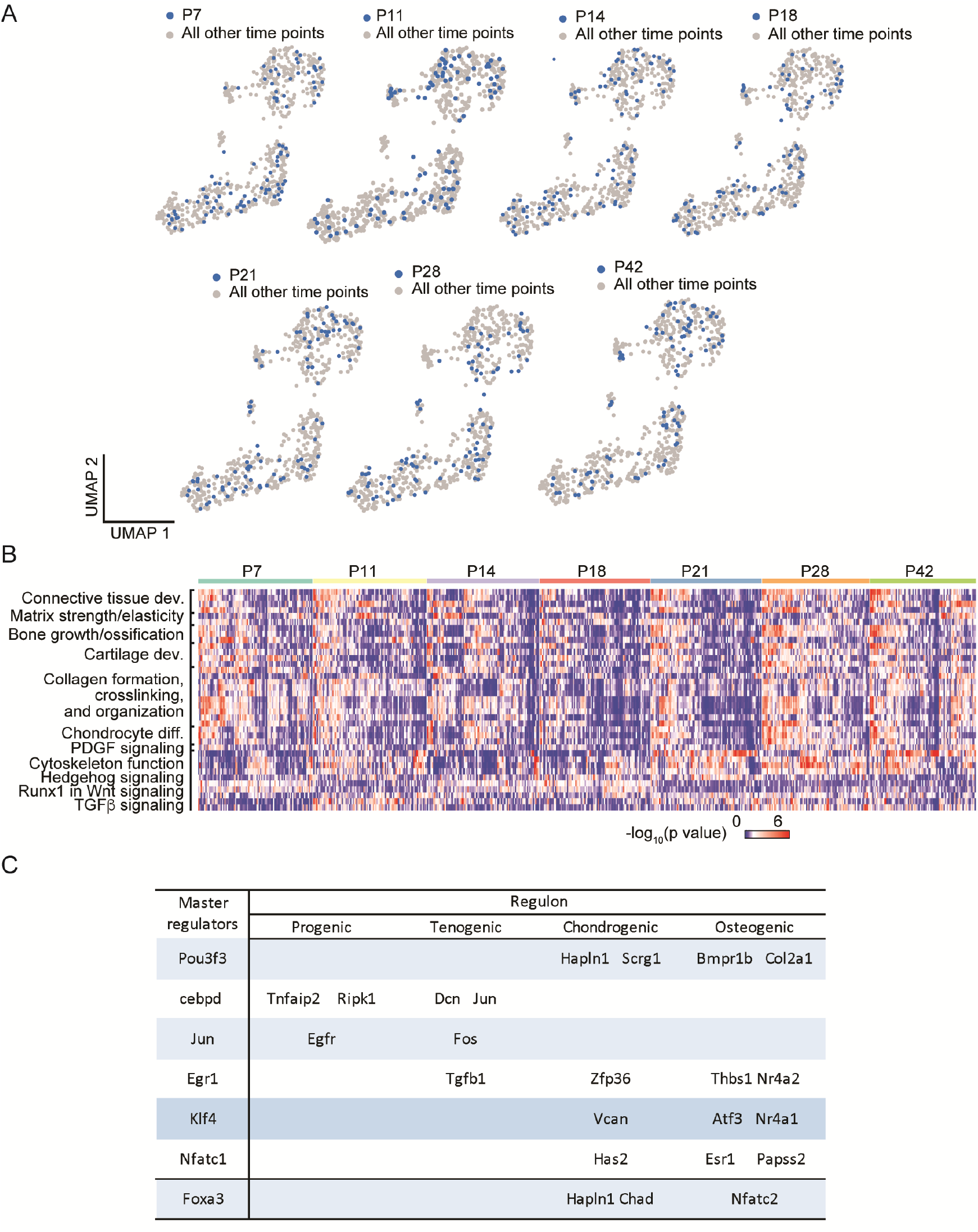
Transcriptional profiling of enthesis Gli1-lineage cells across developmental stages, related to Figure 5 and 6. (A) UMAP plots of Gli-lineage domains at different developmental stages after integration. (B) Heatmap of Gli1-lineage cells revealed the enriched statistical significance of different biological processes for each development stage. (C) Master regulators of Gli1-lineage cells, categorized by gene features, as detected by Scenic analysis.

**Figure S7.**
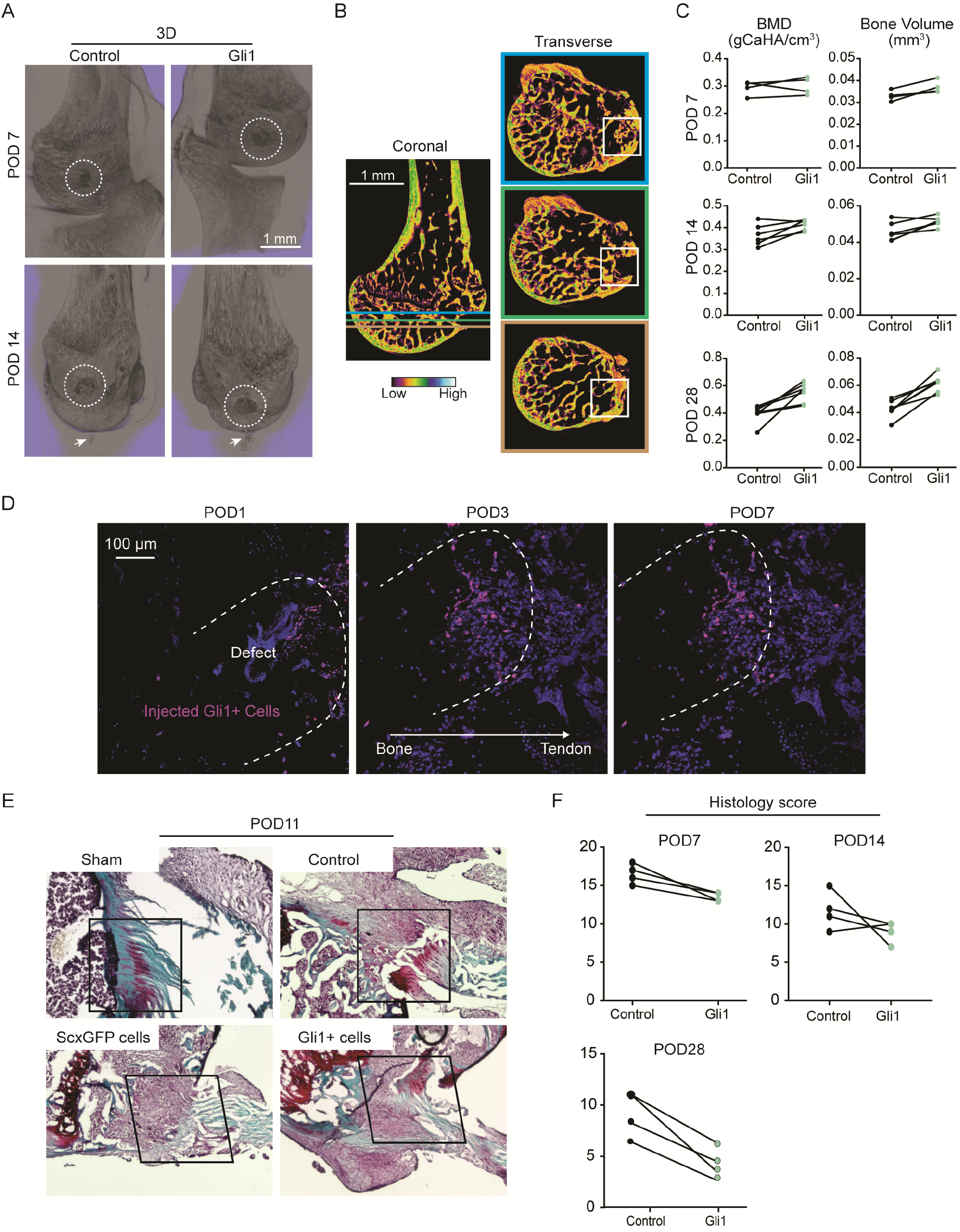
Transplanted Gli1-lineage cells enhance enthesis healing, related to Figure 7. (A) Representative 3D μCT images of humeral head bones with injured entheses at post- operative day 7 (POD7) and POD14. The white dashed circles identify the injured regions and the while arrows point to tissue with heterotopic ossification (HO). Gli1 indicates injuries with transplanted Gli1-lineage cells. Scar bar, 1 mm. (B) Representative coronal images of humeral head bone with three lines corresponding to bottom, middle, and top transverse images, shown on the right. The region of interest used for bone morphometry analysis was selected using a cuboid (projection of white squares) from the top to the bottom of the transverse images. The color bar shows bone density from low to high. (C) Paired comparisons of bone density and bone volume between control (left shoulder, without cell transplantation) and treated (right shoulder, with Gli1-lineage cell transplantation) injuries at POD7, POD14, and POD28. (D) Fluorescent images indicating the retention of transplanted Gli1-lineage cells at the injured enthesis at POD1, POD3, and POD7. Scale bar, 100 μm. The arrow indicates that images are oriented with bone at the left and tendon at the right. (E) Histological images of injured entheses at POD 11 showing more organized extracellular matrix deposition and cell infiltration after transplantation of Gli1-lineage cells, compared to no cell transplantation control, and control transplantation of ScxGFP tenocytes. Scale bar, 100 μm; the black squares highlight the injured enthesis regions; the sections are stained with Safranin O. (F) Paired comparison of histology scores between control (left shoulder, without cell transplantation) and treated tissues (right shoulder, with Gli1-lineage cell transplantation) at POD7, POD14, and POD28.

**Table 1.** Score table for quantifying histological sections of mouse entheses during healing.

|  | **Grade 0** | **Grade 1** | **Grade 2** | **Grade 3** |
| --- | --- | --- | --- | --- |
| **Cellularity** | Hypercellular | Hypercellular, in runs and or increased cell numbers | Hypocellular | Paucicellular |
| **Cell morphology** | Phenotypic gradient in cell morphology; elongated spindle shaped cells near tendon; round cells with lucuna near bone | Heterogeneous distribution of cell morphology; ovoid cell shape near tendon; round cell shape near bone | Homogeneous distribution of cell morphology; some cells near bone show increased roundness and size with decreased amounts of cytoplasm | Homogeneous distribution of cell morphology, with large round nuclei and abundant cytoplasm |
| **Cell orientation** | Cells aligned along collagen fiber direction | Decreased cell alignment in regions near the tendon | Decreased cell alignment across the entire attachment | Random cell alignment |
| **Collagen alignment** | Well-aligned tightly packed collagen fiber bundles | Diminished fiber alignment; separation of individual fiber bundles but with maintenance of overall bundle architecture | Loss of fiber alignment; separation and loss of demarcation of fiber bundles | Randomly aligned fibers; marked separation of fiber bundles with complete loss of architecture |
| **Insertion continuity and maturity** | Proteoglycan staining comparable to normal with clear collagen bundle demarcation | Moderate proteoglycan staining with some loss of collagen bundle demarcation | High proteoglycan staining with clear loss of collagen bundle demarcation | Abundant proteoglycan staining throughout with little to no collagen staining |
| **Bone quality** | Enthesis lamellar and underlying trabecular bone comparable to healthy bone | Formation of thin enthesis lamellar bone and some underlying trabecular bone | Loss of enthesis lamellar bone; maintenance of some underlying trabecular bone | Loss of enthesis lamellar bone and underlying trabecular bone |

Hypocellular: <20 nuclei per field; Hypercellular: 30 nuclei per field. Total score: a score of "0" indicated a healthy enthesis; a score of "18" indicates a pathologic/poorly healed enthesis.

## STAR METHODS

### KEY RESOURCES TABLE

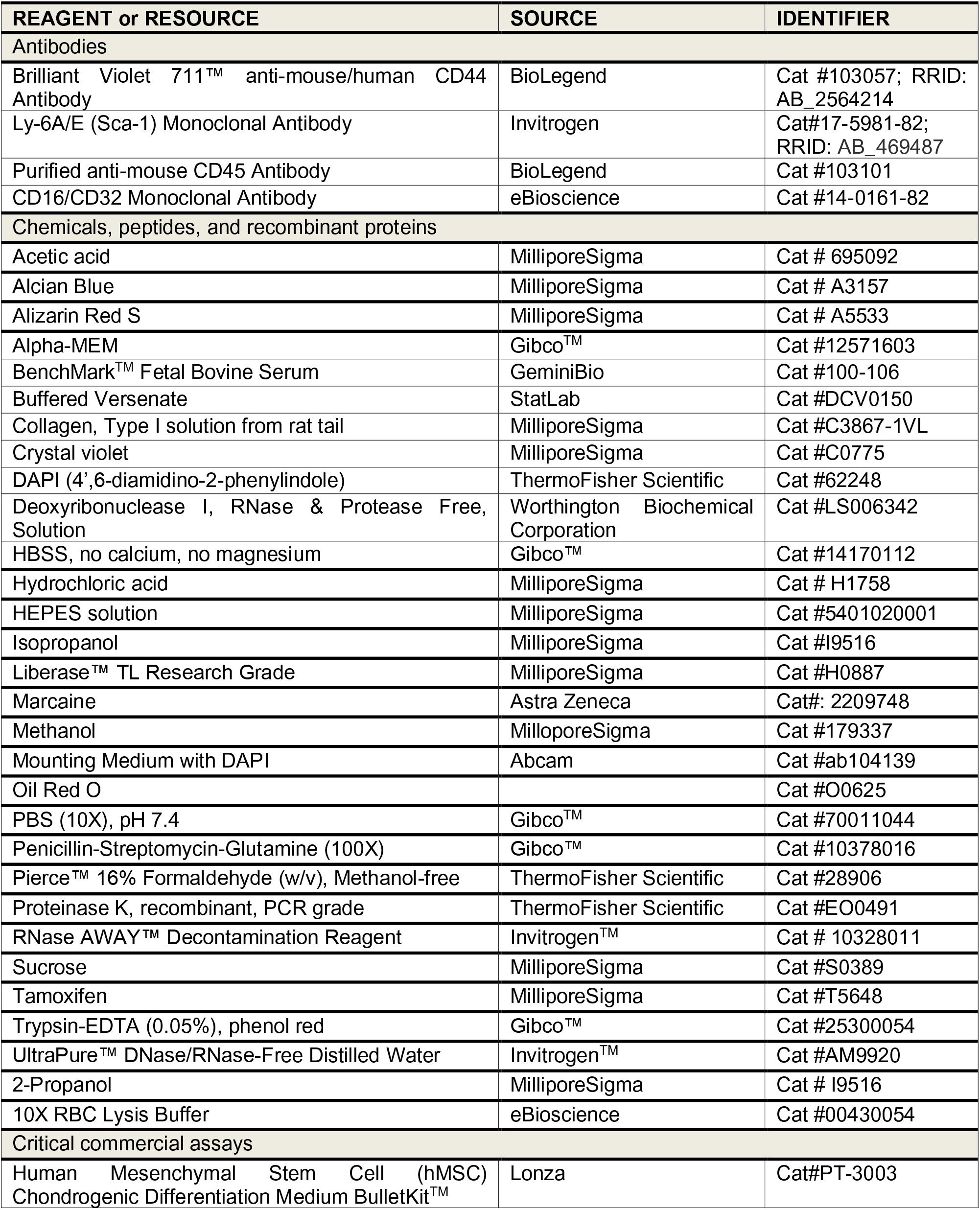
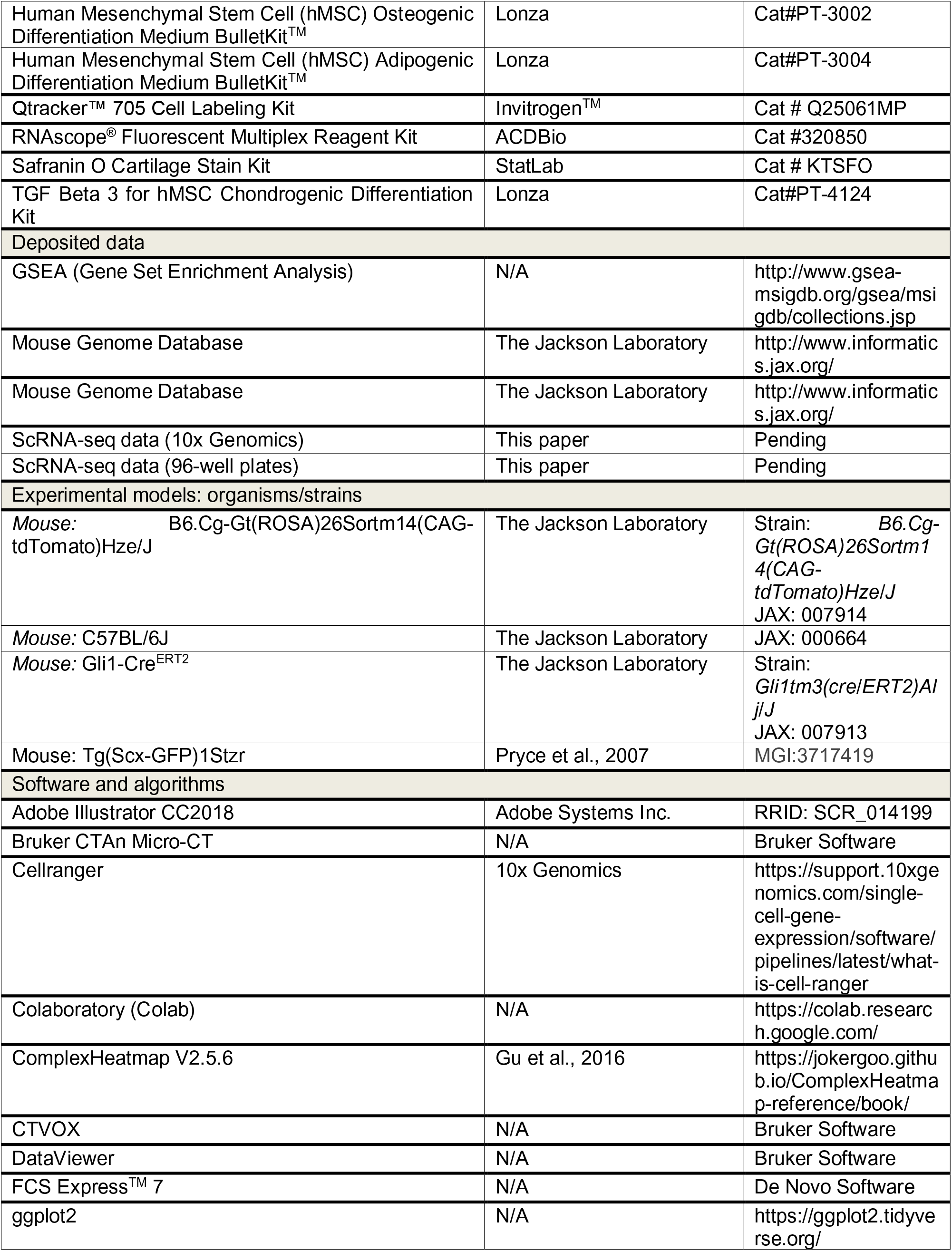
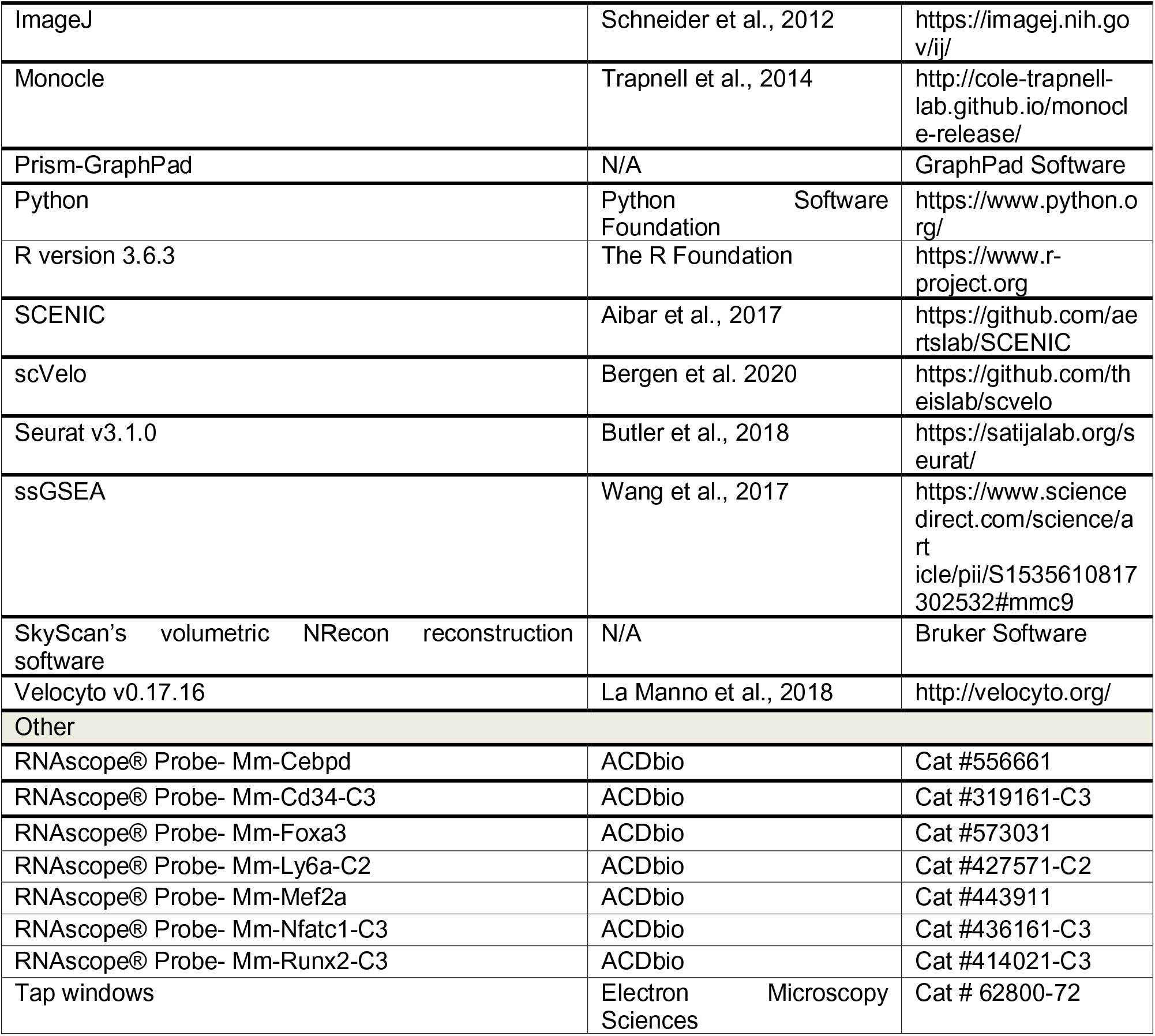

### RESOURCE AVAILABILITY

#### Lead Contact

Further information and requests for resources and reagents should be directed to and will be fulfilled by the Lead Contact, Stavros Thomopoulos.

##### Materials Availability

This work did not generate new reagents or materials.

##### Data and Code Availability

All single-cell datasets created during this study were deposited in the Gene Expression Omnibus (GEO). The GEO access number for the scRNA seq data is GSE182997 and the access token is: gnoxakogdpkxzcj. The data can be accessed with the token at: https://www.ncbi.nlm.nih.gov/geo/query/acc.cgi?acc=GSE182997. All data will be made publicly available at the time of publication.

### EXPERMINETAL MODEL AND SUBJECT DETALS

#### Animal models

All animal procedures were approved by the Columbia University’s institutional Animal Care and Use Committee. Wild-type animals used for 10x Genomics Chromium 3’ Single Cell RNA-Seq analysis and injury models were C57BL/6J mice purchased from Jackson Laboratory. To harvest Gli1-lineage cells for pooled 3’-End scRNA-Seq in 96-well plates, *in vitro* experiment, and cell transplantation for injured models, Gli1Cre^ERT^ mice (Schwartz et al., 2015) were crossed with Ai14 mice. Gli1Cre^ERT^;Ai14 mice were further crossed with ScxGFP mice. Both Gli1Cre^ERT^;Ai14 mice and Gli1Cre^ERT^;Ai14;ScxGFP mice were injected with tamoxifen at P5 and P7 and used in our study. Mice were maintained in a standard cages under a standard 12 h light/dark cycle with *ad libitum* access to food and water.

#### Primary cell culture

Gli1-lineage cells were dissected, dissociated (described below), and sorted by Fluorescence-activated Cell Sorting and then cultured in growth medium containing αMEM medium, 10% fetal bovine serum, and 1% penicillin/streptomycin. Cells were subsequently harvested at passage three for *in vitro* differentiation assays and transplantation in injured mice.

### METHOD DETAILS

#### Single cell isolation for 10x Genomics Chromium

Tendon entheses n=12-16/timepoint for 10 Genomics) from P11, P18, and P56 C57BL/6J mice were dissected under surgical microscope and pooled together (Figure S1B). They were digested in HBSS solution with 300 mg/ml liberase, 20 ng/ml Dnase, 5% FBS, and 100 mg/ml HEPES for 1.5 hours at 37 celsius degree. The samples were filtered through a 40 μm strain, resuspended in the RBC lysis buffer for 5 minutes at room temperature to lyse erythrocytes and stained with Cd45 in 200 μl PBS with 5% FBS for 45 minutes at 4°C to exclude hematopoietic cells. Then, the samples were washed with PBS three times, resuspended in 500 μl PBS with 1 ug/ml DAPI and 5% FBS, and filtered into polypropylene FACS tubes. Single cells without Cd45 and DAPI staining were index sorted by a BD influx Sorter and collected into a microcentrifuge tube for scRNA-seq 10 Genomics experiments.

#### Single cell isolation for scRNA-seq in 96-well plates

For experiments of 3’-End scRNA-Seq in 96-well plates, tendon entheses (n=3/time point) of GliCre^ERT^;Ai14 mice of P7, P11, P18, P21, P28, and P42 were dissected, pooled together, and digested in the aforementioned solution under the same condition discussed above. All cells were stained with 1 ug/ml DAPI and were index sorted by a BD influx Sorter. Single cells without DAPI staining and red fluorescence were collected into 96-well plates, prefilled with lysis buffer for scRNA-Seq analysis.

#### Fluorescence *in situ* hybridization (FISH)

Tendon-bone samples of C57BL/6J mice at P7, P18, and P56 (n=4/timepoint) were fixed in 10% formaldehyde for three hours, permeabilized in 30% sucrose overnight, embedded in OCT compound, and crosectioned into 10 μm-thick tissue slices. Tissue slices were treated with 5 μg/ml Proteinase K for 10 minutes and RNA FISH of these slices was performed using RNAscope FISH techniques according to the manufacturer’s instructions (Barry et al., 2013). To visualize the mRNA spatiotemporal expression pattern, *in situ* hybridization was conducted using the positive and negative probes as control and also the probes against Cebpd, Cd34, Foxa3, Ly6a, Mef2a, Nfatc1, and Runx2. After hybridization, tissue slices were counterstained with DAPI and mounted for images. All FISH image stacks were collected on a Nikon TI Eclipse inverted microscope with a 20X and 60X oil objective for visualization. All imaging parameters of confocal microscope were maintained consistently for FISH. ImageJ software (National Institute of Health) was used to equally perform post-imaging manipulations and quantify percentage of cells with positive staining normalized by the total cell number from histological sections. Maximal projection of image stacks was presented as representative FISH images.

#### Colony formation assay

GliCre^ERT^;Ai14 mice;ScxGFP mice at P11 were injected with tamoxifen injected at P5 and P7. Tendon entheses and partial tendons of three or four GliCre^ERT^;Ai14 mice;ScxGFP mice at P11 were pooled and digested for harvesting cells. All cells were stained with antibody cocktails of Cd44 and Ly6a in PBS for 45 minutes at 4°C, washed three times, and then stained with 1 ug/ml DAPI for sorting. We aimed to sort out three types of cells (n=3-5/each type): Gli1+ Ly6a+ Cd44+ DAPI- enthesis cells (Gli1 progenitors); Gli1+ DAPI- cells (Gli1+ cells); ScxGFP+ Gli1- DAPI- cells (ScxGFP cells). Single cells of three types were sorted in 96-well plates and cultured αMEM medium, 10% fetal bovine serum, and 1% penicillin/streptomycin for 1 month. All were incubated with 0.5% crystal violet staining solution for 20 mins. The wells with more than 100 cells were considered to have colonies. The percentage of colony formation was calculated by normalizing the number of colonies by 96, the total well number.

#### *In vitro* multipotential differentiation assays

Tendon entheses of GliCre^ERT^;Ai14;ScxGFP mice at P11 and P28 (n=4-6/timepoint) with tamoxifen injected at P5 and P7 were pooled together and digested. Gli1+ cells and Gli1- ScxGFP+ cells were collected, cultured in the growth medium, and expanded until passage three for injury treatment. Mouse bone marrow mesenchymal stem cells (BMSCs) were harvested as positive control in parallel. All cells were expanded in αMEM medium, 10% fetal bovine serum, and 1% penicillin/streptomycin. After achieving 80% confluency, cells were trypsinized and separated for adipogenesis, osteogenesis, and chondrogenesis assays.

Cells were seeded on 6-well plates and incubated in culture medium overnight. Cells were then maintained in adipogenic, osteogenic, or chondrogenic differentiation mediums (Lonza) following the manufacturer’s instructions. For checking adipogenesis, cells after culturing in two weeks were fixed in 10% formaldehyde and stained with Oil Red O. For osteogenesis evaluation, cells after maintaining in differentiation medium for 21 days were fixed and stained with 2% Alizarin Red S pH=4.2. For determining chondrogenesis, cells after differentiated for 28 days were stained with Alcian Blue pH=1.2.

#### Cell preparation for transplantation

Tendon entheses of GliCre^ERT^;Ai14;ScxGFP mice at P1. were pooled together and digested for harvesting Gli1+ cells. Gli1+ cells were collected, cultured in the growth medium, and expanded until passage three for injury treatment. To check the retention of Gli1+ cells at post-operative day (POD) 1, 3, and 7, cells were released from culturing dishes, incubated in Qtracker 705 Cell Labeling Kit for 1 hour following its protocol and washed twice. Finally, all cells were suspended in sterile 4mg/ml Collagen type I solution from rat tail at the concentration of 2X10^7^ cells/ml.

#### Needle punch enthesis injury and cell transplantation

Enthesis injury protocols were adapted from our previous study (Schwartz et al., 2017). After injection with buprenorphine SR and then isoflurane anesthesia, nine to ten-week-old C57BL/6J mice (n=6-8 with females and males for each group of post-operative day sacrifice) were placed in a lateral decubitus position. The upper limb was externally rotated to bring the forearm in supination and secured. An incision was made in the skin to visualize the deltoid. The deltoid was cut to expose the supraspinatus enthesis and humerus head. For the injured right shoulder treated with Gli1-lineage cells, 1X10^6^ cells in 50 μl Collagen type I solution and pumped into a syringe with a 28G needle. The needle was inserted into the tendon enthesis (close to the humeral head) to create a punch defect until the enthesis was completely bisected into the marrow cavity. The cells were gradually released from the syringe to fill the punched hole. For the contralateral left shoulder as control, only the punch injury was created. 5-0 PROLENE suture was used to ligate the deltoid back over the humerus and close the skin. After injury, mice experienced free cage activity and euthanized after 1, 2, and four weeks.

#### Single cell RNA sequencing

For 10x Genomics Chromium 3’ solution, the cells were centrifuged at 500g for 5 mins, counted by a hemocytometer, and resuspended in PBS with 10% FBS. Approximately 5,000 cells were captured for each sample. The sorted cells were loaded into 10x Chromium Controller using the Chromium Single Cell 3’ v3 reagents and prepared as droplets for lysis and reverse transcription. The resulting pooled, 3’-end libraries were sequenced on Illumina® NovaSeq™ 6000 Sequencer.

For scRNA-Seq in 96-well plates, 96-well plates with cells in lysis buffer were spun down and processed with automated liquid handling robot. Adapter-linked oligo(dT) primers including both cell- and molecule- specific barcodes were used to complete template-switching reverse transcription. Pooled 3’-end sequencing libraries were finally sequenced on an IIlumina NextSeq 500.

#### Single cell RNA sequencing data analysis

Cell Ranger was used to demultiplex raw data for creating Fastq files. Following alignment and filtering, gene-level unique molecular identifier counts (UMIs) and gene expression matrix were obtained for downstream analysis. R package Seurat was used as a first analytical tool to load data and build Seurat objects, which were processed for controlling data quality, filtering, clustering, and data visualization, and examining differential expression analysis. Specifically, cells with genes expressed less than 2 cells, cells<1000 UMIs with >20% UMIs mapped to mitochondrial genes, and <200 genes were removed. After normalization of cells by the total UMI read counts, unsupervised shared nearest neighbor clustering (SNN) with an adjusted resolution were performed and displayed in Umap. Clusters were subset and identified by representative marker genes related with, for example, tenogenesis and chondrogenesis. Datasets at different timepoints were integrated via canonical correlation analysis (CCA) and combined into a uniform Umap Alta. Shared patterns of transcriptional profile were highlighted by CCA analysis and shown by heatmap and dot plots.

#### Gene regulatory network analysis

For single cell regulatory network analysis, the SCENIC package (V1.2.2) was used with default settings to identify transcription networks. First, GENIE3 module identified target genes that were significantly co- expressed with a certain transcription factor. Next, RcisTarget module predicted the target genes by screening the enriched cis-regulatory motifs of candidate transcription factors. Lastly, AUCell algorithm scored the activity of each regulon. Regulon activities to show the master regulators of each single cell were plotted by heatmap.

#### Gene set enrichment analysis

Single sample gene set enrichment analysis (ssGSEA), which identified the enriched gene sets at a single cell level, was as previously described (Wang et al., 2017). In brief, we computed enrichment scores of a certain gene set for both the experimental dataset and a random permutated dataset of 1000 cells. Normalized gene expression matrix from Seurat analysis and gene sets of interest (from GSEA Molecular Signatures Database v7.2) were used as inputs. *p* values of each gene set were computed by comparing our dataset to the permutated dataset. The heatmaps showed –log10(*p* values) of enriched gene sets using the ComplexHeatmap package (Gu et al., 2016).

#### Cell lineage trajectory analysis

Following the instructed tutorial of Monocle 3 package with the default settings, detailed pseudotime for tendon enthesis resident cells over different time points (Trapnell et al., 2014). Cell clusters were also arranged along the pseudotime to represent the lineage trajectory. For RNA velocity, a loom file of spliced and unspliced mRNAs from aligned bam file using Velocyto package (V0.17.16) in Python (V3.6) was created. Then RNA velocity was computed based on both steady state and dynamic models using scVelo package (Bergen et al., 2020). The scVelo algorithm used raw read counts from sequencing and was insensitive to library size normalization.

#### Microcomputed tomography

Tendon-bone samples from injury mice were dissected and fixed in 10% formaldehyde for bone morphometry analysis (Fang et al., 2020). Microcomputed tomography (μCT, Bruker Skyscan 1272) with an energy of 55 kilovolt peaks, an intensity of 145 μA, and a standard resolution of 5 μm was used to scan samples. After image reconstruction, region of interest around the injury region was selected by a rectangle with a fixed dimension (Figure S7B). Bone volume (mm^3^), BMD, Tb.Th in the region of interest were measured directly with CTAn (Bruker). For 3D visualization, the reconstructed images were imported into CTVOX (Bruker).

#### Histology

Tendon-bone samples after μCT scanning were dissected and fixed 10% formaldehyde overnight. Following this, tendon-bone samples were decalcified in Buffered Versenate for two weeks. Subsequently, the samples were embedded in optimal cutting temperature (OCT) compound, cyrosectioned into 10 μm- thick slices, and stained with safranin O according to the standard protocol. All images were collected on a Nikon TI Eclipse inverted microscope for visualization and semi-quantified according to the table. Similarly, tissue sections of injured enthesis at POD7 to track the retention of Gli1-lineage cells were prepared. Tissue slices were counterstained with DAPI and mounted for imaging by using a Nikon TI Eclipse inverted microscope with a 20X and 60X oil objective discussed above.

### QUANTIFICATION AND STATISTICAL ANALYSIS

Graphpad Prism 7 was used to conduct statistical analysis. Results are shown as mean±SD. For every experiment, both male and female mice from at least two independent litters were used. All data analysis were conducted blindly. When comparison was applied between two groups, an unpaired or paired (when appropriate) Student’s t test was used. When comparison was applied among multiple groups or two factors, ANOVA was performed with post hoc Tukey correction. 0.05<#p<0.1, *p < 0.05; **p < 0.01; ***p < 0.001; ****p < 0.0001 are shown to indicate statistically significantly difference in figures. The exact numbers of repeated sample for each experiment are revealed in the corresponding figure legends.

## References

Aibar, S., González-Blas, C.B., Moerman, T., Imrichova, H., Hulselmans, G., Rambow, F., Marine, J.C., Geurts, P., Aerts, J., and van den Oord, J. (2017). SCENIC: single-cell regulatory network inference and clustering. Nature methods 14, 1083–1086.

Akiyama, H., Chaboissier, M.C., Martin, J.F., Schedl, A., and de Crombrugghe, B. (2002). The transcription factor Sox9 has essential roles in successive steps of the chondrocyte differentiation pathway and is required for expression of Sox5 and Sox6. Genes & development 16, 2813–2828.

Alvarez-Dolado, M., Pardal, R., Garcia-Verdugo, J.M., Fike, J.R., Lee, H.O., Pfeffer, K., Lois, C., Morrison, S.J., and Alvarez-Buylla, A. (2003). Fusion of bone-marrow-derived cells with Purkinje neurons, cardiomyocytes and hepatocytes. Nature 425, 968–973.

Amakye, D., Jagani, Z., and Dorsch, M. (2013). Unraveling the therapeutic potential of the Hedgehog pathway in cancer. Nature medicine 19, 1410.

Barry, E.R., Morikawa, T., Butler, B.L., Shrestha, K., de La Rosa, R., Yan, K.S., Fuchs, C.S., Magness, S.T., Smits, R., and Ogino, S. (2013). Restriction of intestinal stem cell expansion and the regenerative response by YAP. Nature 493, 106–110.

Bergen, V., Lange, M., Peidli, S., Wolf, F.A., and Theis, F.J. (2020). Generalizing RNA velocity to transient cell states through dynamical modeling. Nature biotechnology 38, 1408–1414.

Best, K.T., Nichols, A.E., Knapp, E., Hammert, W.C., Ketonis, C., Jonason, J.H., Awad, H.A., and Loiselle, A. (2020). NF-κB activation persists into the remodeling phase of tendon healing and promotes myofibroblast survival. Science signaling 13, 658.

Bi, Y., Ehirchiou, D., Kilts, T.M., Inkson, C.A., Embree, M.C., Sonoyama, W., Li, L., Leet, A.I., Seo, B.M., and Zhang, L. (2007). Identification of tendon stem/progenitor cells and the role of the extracellular matrix in their niche. Nature medicine 13, 1219–1227.

Blitz, E., Sharir, A., Akiyama, H., and Zelzer, E. (2013). Tendon-bone attachment unit is formed modularly by a distinct pool of Scx-and Sox9-positive progenitors. Development 140, 2680–2690.

Carpenter, J., Thomopoulos, S., Flanagan, C., DeBano, C., and Soslowsky, L. (1998). Rotator cuff defect healing: a biomechanical and histologic analysis in an animal model. Journal of Shoulder and Elbow Surgery 7, 599–605.

Chan, C.K., Gulati, G.S., Sinha, R., Tompkins, J.V., Lopez, M., Carter, A.C., Ransom, R.C., Reinisch, A., Wearda, T., and Murphy, M. (2018). Identification of the human skeletal stem cell. Cell 175, 43–56. e21.

Chen, Z., Lu, Y., Zhang, K., Xiao, Y., Lu, J., and Fan, R. (2019). Multiplexed, sequential secretion analysis of the same single cells reveals distinct effector response dynamics dependent on the initial basal state. Advanced Science 6, 1801361.

De Micheli, A.J., Laurilliard, E.J., Heinke, C.L., Ravichandran, H., Fraczek, P., Soueid-Baumgarten, S., De Vlaminck, I., Elemento, O., and Cosgrove, B.D. (2020). Single-cell analysis of the muscle stem cell hierarchy identifies heterotypic communication signals involved in skeletal muscle regeneration. Cell reports 30, 3583–3595. e3585.

Degirmenci, B., Valenta, T., Dimitrieva, S., Hausmann, G., and Basler, K. (2018). GLI1-expressing mesenchymal cells form the essential Wnt-secreting niche for colon stem cells. Nature 558, 449–453.

Derwin, K.A., Galatz, L.M., Ratcliffe, A., and Thomopoulos, S. (2018). Enthesis repair: challenges and opportunities for effective tendon-to-bone healing. The Journal of bone and joint surgery American volume 100, e109.

Fang, F., and Lake, S.P. (2017). Multiscale mechanical evaluation of human supraspinatus tendon under shear loading after glycosaminoglycan reduction. Journal of biomechanical engineering 139, 7.

Fang, F., and Lake, S.P. (2015). Multiscale strain analysis of tendon subjected to shear and compression demonstrates strain attenuation, fiber sliding, and reorganization. Journal of orthopaedic research 33, 1704–1712.

Fang, F., and Lake, S.P. (2016). Multiscale mechanical integrity of human supraspinatus tendon in shear after elastin depletion. Journal of the mechanical behavior of biomedical materials 63, 443–455.

Fang, F., Schwartz, A.G., Moore, E.R., Sup, M.E., and Thomopoulos, S. (2020). Primary cilia as the nexus of biophysical and hedgehog signaling at the tendon enthesis. Science advances 6, eabc1799.

Felsenthal, N., Rubin, S., Stern, T., Krief, S., Pal, D., Pryce, B.A., Schweitzer, R., and Zelzer, E. (2018). Development of migrating tendon-bone attachments involves replacement of progenitor populations. Development 145, dev165381.

Feng, H., Xing, W., Han, Y., Sun, J., Kong, M., Gao, B., Yang, Y., Yin, Z., Chen, X., and Zhao, Y. (2020). Tendon- derived cathepsin K–expressing progenitor cells activate Hedgehog signaling to drive heterotopic ossification. The Journal of Clinical Investigation 130.

Galatz, L.M., Ball, C.M., Teefey, S.A., Middleton, W.D., and Yamaguchi, K. (2004). The outcome and repair integrity of completely arthroscopically repaired large and massive rotator cuff tears. JBJS 86, 219–224.

Genin, G.M., Kent, A., Birman, V., Wopenka, B., Pasteris, J.D., Marquez, P.J., and Thomopoulos, S. (2009). Functional grading of mineral and collagen in the attachment of tendon to bone. Biophysical journal 97, 976–985.

Gracey, E., Burssens, A., Cambré, I., Schett, G., Lories, R., McInnes, I.B., Asahara, H., and Elewaut, D. (2020). Tendon and ligament mechanical loading in the pathogenesis of inflammatory arthritis. Nature reviews rheumatology 16, 193–207.

Graham, G., Wright, E., Hewick, R., Wolpe, S., Wilkie, N., Donaldson, D., Lorimore, S., and Pragnell, I. (1990). Identification and characterization of an inhibitor of haemopoietic stem cell proliferation. Nature 344, 442–444.

Gu, Z., Eils, R., and Schlesner, M. (2016). Complex heatmaps reveal patterns and correlations in multidimensional genomic data. Bioinformatics 32, 2847–2849.

Harada, S., and Rodan, G.A. (2003). Control of osteoblast function and regulation of bone mass. Nature 423, 349–355.

Harryman, D.T., Hettrich, C.M., Smith, K.L., Campbell, B., Sidles, J.A., and Matsen III, F.A. (2003). A prospective multipractice investigation of patients with full-thickness rotator cuff tears: the importance of comorbidities, practice, and other covariables on self-assessed shoulder function and health status. JBJS 85, 690–696.

Harvey, T., Flamenco, S., and Fan, C.M. (2019a). A Tppp3+ Pdgfra+ tendon stem cell population contributes to regeneration and reveals a shared role for PDGF signalling in regeneration and fibrosis. Nature cell biology 21, 1490–1503.

Jiang, Y., Jahagirdar, B.N., Reinhardt, R.L., Schwartz, R.E., Keene, C.D., Ortiz-Gonzalez, X.R., Reyes, M., Lenvik, T., Lund, T., and Blackstad, M. (2002). Pluripotency of mesenchymal stem cells derived from adult marrow. Nature 418, 41–49.

Khan, S.K., Yadav, P.S., Elliott, G., Hu, D.Z., Xu, R., and Yang, Y. (2018). Induced GnasR201H expression from the endogenous Gnas locus causes fibrous dysplasia by up-regulating Wnt/β-catenin signaling. Proceedings of the National Academy of Sciences 115, E418–E427.

Kramann, R., Goettsch, C., Wongboonsin, J., Iwata, H., Schneider, R.K., Kuppe, C., Kaesler, N., Chang-Panesso, M., Machado, F.G., and Gratwohl, S.J. (2016). Adventitial MSC-like cells are progenitors of vascular smooth muscle cells and drive vascular calcification in chronic kidney disease. Cell stem cell 19, 628–642.

Kramann, R., Schneider, R.K., DiRocco, D.P., Machado, F., Fleig, S., Bondzie, P.A., Henderson, J.M., Ebert, B.L., and Humphreys, B.D. (2015). Perivascular Gli1+ progenitors are key contributors to injury-induced organ fibrosis. cell stem cell 16, 51–66.

Kronenberg, H.M. (2003). Developmental regulation of the growth plate. Nature 423, 332–336.

Leupin, O., Kramer, I., Collette, N.M., Loots, G.G., Natt, F., Kneissel, M., and Keller, H. (2007). Control of the SOST bone enhancer by PTH using MEF2 transcription factors. Journal of bone and mineral research 22, 1957–1967.

Lindvall, O., Kokaia, Z., and Martinez-Serrano, A.J. (2004). Stem cell therapy for human neurodegenerative disorders–how to make it work. Nature medicine 10, S42–S50.

Liu, H., Xu, J., and Jiang, R. (2019). Mkx-Deficient Mice Exhibit Hedgehog Signaling–Dependent Ectopic Ossification in the Achilles Tendons. Journal of Bone and Mineral Research 34, 557–569.

Long, F., and Ornitz, D.M. (2013). Development of the endochondral skeleton. Cold spring harbor perspectives in biology 5, a008334.

Lu, H.H., and Thomopoulos, S. (2013). Functional attachment of soft tissues to bone: development, healing, and tissue engineering. Annual review of biomedical engineering 15, 201–226.

Men, Y., Wang, Y., Yi, Y., Jing, D., Luo, W., Shen, B., Stenberg, W., Chai, Y., Ge, W.P., and Feng, J.Q.. (2020). Gli1+ periodontium stem cells are regulated by osteocytes and occlusal force. Developmental cell 54, 639–654. e636.

Millar, N.L., Silbernagel, K.G., Thorborg, K., Kirwan, P.D., Galatz, L.M., Abrams, G.D., Murrell, G.A., McInnes, I.B., and Rodeo, S. (2021). Tendinopathy. 7, 1–21.

Novince, C.M., Michalski, M.N., Koh, A.J., Sinder, B.P., Entezami, P., Eber, M.R., Pettway, G.J., Rosol, T.J., Wronski, T.J., and Kozloff, K.M. (2012). Proteoglycan 4: a dynamic regulator of skeletogenesis and parathyroid hormone skeletal anabolism. Journal of Bone and Mineral Research 27, 11–25.

Schett, G., Lories, R.J., D’Agostino, M.A., Elewaut, D., Kirkham, B., Soriano, E.R., and McGonagle, D. (2017). Enthesitis: from pathophysiology to treatment. Nature reviews rheumatology 13, 731.

Schneider, R.K., Mullally, A., Dugourd, A., Peisker, F., Hoogenboezem, R., Van Strien, P.M., Bindels, E.M., Heckl, D., Büsche, G., and Fleck, D. (2017). Gli1+ mesenchymal stromal cells are a key driver of bone marrow fibrosis and an important cellular therapeutic target. Cell stem cell 20, 785–800. e788.

Schwartz, A.G., Galatz, L.M., and Thomopoulos, S. (2017). Enthesis regeneration: a role for Gli1+ progenitor cells. Development 144, 1159–1164.

Schwartz, A.G., Long, F., and Thomopoulos, S. (2015). Enthesis fibrocartilage cells originate from a population of Hedgehog-responsive cells modulated by the loading environment. Development 142, 196–206.

Schwartz, A.G., Pasteris, J.D., Genin, G.M., Daulton, T.L., and Thomopoulos, S. (2012). Mineral distributions at the developing tendon enthesis. PloS one 7, e48630.

Shi, Y., He, G., Lee, W.C., McKenzie, J.A., Silva, M.J., and Long, F.X. (2017). Gli1 identifies osteogenic progenitors for bone formation and fracture repair. Nature communications 8, 1–12.

Sugimoto, Y., Takimoto, A., Akiyama, H., Kist, R., Scherer, G., Nakamura, T., Hiraki, Y., and Shukunami, C. (2013). Scx+/Sox9+ progenitors contribute to the establishment of the junction between cartilage and tendon/ligament. Development 140, 2280–2288.

Thomopoulos, S., Genin, G.M., and Galatz, L.M. (2010). The development and morphogenesis of the tendon-to- bone insertion What development can teach us about healing. Journal of musculoskeletal & neuronal interactions 10, 35.

Thomopoulos, S., Williams, G.R., Gimbel, J.A., Favata, M., and Soslowsky, L.J. (2003). Variation of biomechanical, structural, and compositional properties along the tendon to bone insertion site. Journal of orthopaedic research 21, 413–419.

Tikhonova, A.N., Dolgalev, I., Hu, H., Sivaraj, K.K., Hoxha, E., Cuesta-Domínguez, Á., Pinho, S., Akhmetzyanova, I., Gao, J., and Witkowski, M. (2019). The bone marrow microenvironment at single-cell resolution. Nature 569, 222–228.

Trapnell, C., Cacchiarelli, D., Grimsby, J., Pokharel, P., Li, S., Morse, M., Lennon, N.J., Livak, K.J., Mikkelsen, T.S., and Rinn, J.L. (2014a). The dynamics and regulators of cell fate decisions are revealed by pseudotemporal ordering of single cells. Nature biotechnology 32, 381.

van den Brink, S.C., Sage, F., Vértesy, Á., Spanjaard, B., Peterson-Maduro, J., Baron, C.S., Robin, C., and Van Oudenaarden, A. (2017). Single-cell sequencing reveals dissociation-induced gene expression in tissue subpopulations. Nature methods 14, 935–936.

Wang, Q., Hu, B., Hu, X., Kim, H., Squatrito, M., Scarpace, L., deCarvalho, A.C., Lyu, S., Li, P., and Li, Y.J.C.c. (2017a). Tumor evolution of glioma-intrinsic gene expression subtypes associates with immunological changes in the microenvironment. Cancel cell 32, 42–56. e46.

Wang, Y., Zhang, X., Huang, H., Xia, Y., Yao, Y., Mak, A.F.-T., Yung, P.S.-H., Chan, K.-M., Wang, L., and Zhang, C. (2017b). Osteocalcin expressing cells from tendon sheaths in mice contribute to tendon repair by activating Hedgehog signaling. Elife 6, e30474.

Zhao, H., Feng, J., Seidel, K., Shi, S., Klein, O., Sharpe, P., and Chai, Y. (2014). Secretion of shh by a neurovascular bundle niche supports mesenchymal stem cell homeostasis in the adult mouse incisor. Cell stem cell 14, 160–173.

Zhong, L., Yao, L., Tower, R.J., Wei, Y., Miao, Z., Park, J., Shrestha, R., Wang, L., Yu, W., and Holdreith, N.J.E. (2020). Single cell transcriptomics identifies a unique adipose lineage cell population that regulates bone marrow environment. Elife 9, e54695.

